# Remote Phenotyping Strategies for Sunflower Flowering Assessments Using Deep Learning Approaches

**DOI:** 10.1101/2025.04.03.646664

**Authors:** Leonardo Volpato, Sara Tirado Tolosa, Chaitanyam Potnuru, Jason Jaacks, Jamie Layton

## Abstract

The flowering date of sunflowers is a crucial trait that significantly influences crop management practices and product placement. Traditional ground methods for data collection are labor-intensive and subjective, requiring field scientists to manually estimate and record data in the field. This trait can be measured by counting the number of days from planting until 50% of plants in each research plot have reached flowering at R5 developmental growth stage. However, this method is time-consuming and may overlook valuable information related to flowering rates and duration. Flowering time of sunflower also can be approximated by counting the number of heads (flowers) across multiple dates. We propose a method for rapidly counting sunflower heads to model flower counts over time and estimate flowering time using RGB images acquired by Unmanned Aerial Vehicles (UAVs). The method developed employs a deep learning model trained to detect sunflower heads from UAV imagery and modeling these counts over time using a logistic function to estimate the 50% flowering date. The experimental results obtained from this method enabled estimation of the flowering date with a high correlation to ground measurements (*r* > 0.91). Significantly, this approach not only reduces labor but also improves the precision of data collection. Moreover, an increase of 6% in heritability across trials, compared to traditional methods, suggests that our approach contributes to a deeper genetic understanding of flowering dynamics. This includes enhanced insights into the timing and rates of flowering, essential for optimizing breeding strategies and understanding genetic responses to environmental conditions. This innovative approach offers a promising avenue for enhancing the efficiency and accuracy of sunflower phenotyping.

## 1. INTRODUCTION

Sunflowers (*Helianthus annuus*) are a globally significant crop valued for their seeds and oil (Puttha et al., 2023). Sunflower delivers ornamental value, medicinal use and health-promoting properties which provide a valuable source of nutrition and income for millions of people around the world (S. Guo et al., 2017). The efficiency of sunflower cultivation directly impacts global food production and economic stability in the agricultural sector. Sunflower yield and oil production can be significantly affected by flowering dates, which are controlled by genetic and environmental factors.

The flowering date of sunflowers is a critical determinant in crop management and yield optimization (Craufurd & Wheeler, 2009). Flowering date influences key decisions in the agricultural process, from planting schedules to harvest timing. Recording flowering data has helped researchers to study flowering patterns for several important crops such as maize (*Zea mays* L.) (Fan et al., 2022; Prasad et al., 2018), rice (*Oryza sativa* L.) (Lian et al., 2024; Sohail, 2023), wheat (*Triticum aestivum* L.) (P. Hu et al., 2021, 2021), barley (*Hordeum vulgare* L.) (Ibrahim et al., 2018; Kupke et al., 2022), and sorghum (*Sorghum bicolor*) (Casto et al., 2019). Characterization of flowering patterns facilitates genetic and physiological studies of angiosperms, such as cereal crops, and contributes to breeding novel cultivars for optimal yield and environmental adaptability (Hill & Li, 2016).

Traditionally, the estimation of flowering date in sunflower cultivation is conducted through labor-intensive, ground-based methods. Sunflower plots are monitored regularly by human evaluators who note characteristics such as the number of days after planting until the appearance of flowers in 50% of the plants within a given plot. This process, characterized by manual observation and recording, presents numerous challenges such as time consumption, subjectivity, and potential for data inaccuracy. These manual observations are also subjective and error-prone, as different observers may perceive the growth stage of the same plot differently, introducing variability and potential inaccuracies in the data. In addition, human evaluators often use a categorical scoring system to assess flowering stages (e.g., estimating when 50% of plants in a plot have flower heads starting to open), so that the time duration between flowering stages can be calculated. However, due to time constraints, plots are rarely visited daily, requiring evaluators to estimate data for non-visited days. To effectively reduce the acquisition cost and improve efficiency in data collection, we propose an automatic method for measuring sunflower flowering dates within a breeding program.

Recent advancements in agricultural technology have introduced Unmanned Aerial Vehicles (UAVs), generically referred to as drones, as a potential solution to traditional agricultural challenges providing an efficient and cost-effective crop monitoring solution. Remote sensing (RS) data from drones equipped with advanced imaging capabilities offer a new perspective on crop monitoring and has helped farmers better identify their crop growth periods, assess their crop health, and make informed decisions about crop management (Gano et al., 2024; Iost Filho et al., 2020; Sweet et al., 2022; H. Wang et al., 2023). Past studies have used this technology to identify growth periods and perform crop monitoring for sunflower breeding fields (Z. Song et al., 2023). Investigators have used drones to detect sunflower lodging in low-density fields. Studies have integrated deep learning (DL) techniques with drone-based imagery to offer a transformative approach to agricultural practices, enabling more precise and sophisticated analysis of sunflower growth patterns and health indicators (G. Li et al., 2021; Z. Song et al., 2020). These techniques, which leverage complex algorithms and large datasets, can significantly optimize crop management and yield predictions by enhancing the accuracy and efficiency of detecting and interpreting various growth stages and stress factors in sunflowers.

Advances in HTP (High-Throughput Phenotyping) and breakthroughs in DL enable the rapid characterization of phenotypes like flowering. Several studies demonstrated the use of deep convolutional neural networks (CNNs) and meta-models (e.g., Faster RCNN - Ren et al., 2017) to detect and count plant features to support breeding programs (Jiang & Li, 2020; Khaki et al., 2022). Combining DL with CNN vastly outperformed the traditional machine learning systems in many applications (Murphy et al., 2024; Selmy et al., 2024; Tong et al., 2022). In sunflower, DL has been used to provide sunflower lodging assessments and the sunflower growth period (G. Li et al., 2021; Z. Song et al., 2020, 2023).

Various CNN architectures have been developed for computer vision tasks such as image classification, object detection, and semantic and instance segmentation (Adegun, 2023; Alzubaidi, 2021). Depending on the data format, there are several types of CNN models that can be applied for the detection and counting of plants and plant organs to perform plant development characterization (Lastname et al., 2020; Neupane et al., n.d.; Tong et al., 2022). In regression-based counting strategies, the DL-based density estimation model estimates a density map from an image that represents the distribution and density of the target objects (e.g., flowers) within the image (Peng et al., 2022). The total count of these objects is then derived by summing up the values in the density map. Semantic segmentation also can provide insights into object detection by applying masks for target objects with the same semantic meaning (e.g., all flowers in an image) (Manakitsa et al., 2024). This approach can be used to perform flower counting by leveraging morphological filtering and heuristic rules. Semantic segmentation using DL methods has provided an effective method to perform stand count in several crops using RS data (Lu et al., 2024; Volpato et al., 2023; Y.-H. Wang & Su, 2022). However, the success of the method relies on pre-adjusted parameters to filter the background (non-target pixels), which is frequently not uniform across environments (Emek Soylu et al., 2023; Shaaban et al., 2020). To overcome the limitations of semantic segmentation, a U-net based density estimation model has been proposed for flower counting in several crops (Hobbs et al., 2020; Tao et al., 2022). This model offers significant advancement by accurately identifying and counting target objects on a pixel-by-pixel basis. Furthermore, it demonstrates robustness against varying environmental conditions, which is crucial for reliable performance in diverse agricultural settings. DL-based solutions have been extensively tested for flower detection and counting, demonstrating high accuracy (R2 > 0.92) as compared with ground-truth (GT) measurements for field crops such as wheat (Xu et al., 2023; Ye et al., 2023), cotton (Jiang et al., 2020), corn (Fan et al., 2022), sorghum (H. Li et al., 2022) and rice (X. Wang et al., 2022). Therefore, these studies have demonstrated that aerial images can provide efficient data to feed DL methods to perform object detection based on either a single image or temporal data set. However, limited studies have been found to estimate sunflower growth stage dates. Furthermore, flower detection and counting in sunflower crops can be monitored across time to capture several data points in a single experimental unit (plant or plot) during the entire flowering period to characterize key flowering patterns. Time series data can be modeled by using a simple linear model or a nonlinear model, such as a logistic function, to link the model parameters with the characteristic phenological process (Diao & Li, 2022; Lever et al., 2016). Logistic regression curves have been widely utilized to model vegetation index satellite time series data with vegetation phenological dynamics (Roy & Yan, 2020). In soybeans, a logistic growth function has been deployed to model longitudinal data of canopy coverage through the season to yield improvement (Xavier et al., 2017). Therefore, using time-series data obtained from aerial imagery of drones, logistic regression can effectively model the probability of a plant reaching a certain developmental stage, such as flowering, at different time points.

The integration of logistic regression models and CNN-based image analysis can improve precision agriculture practices. By accurately predicting flowering times, farmers can optimize irrigation schedules, pest management, and harvesting periods, leading to improved yield and resource efficiency. Precise flowering data can be linked with weather data as well, to identify potential stressors during flowering that might signal yield penalty from excess heat or water deficit. Additionally, these techniques can aid in breeding programs by allowing for the selection of cultivars with desirable phenotypes, such as early or late flowering, adapted to specific environmental conditions. There is a lack of comprehensive studies utilizing UAV technologies for the precise estimation of sunflower flowering dates. Our research aims to fill this gap by leveraging UAV-based imagery and DL models to accurately predict the 50% flowering date in sunflowers. Our study successfully demonstrates a high correlation between UAV-derived estimates of flowering dates and traditional human-derived GT measurements, with additional insights into flowering dynamics in sunflower, such as flowering duration and homogeneity. The deployed approach proved to be cost-effective and accurate for assessing sunflower 50% flowering date, which can be used to better manage sunflower production and enhance product recommendations and development.

## 2. MATERIALS AND METHODS

### 2.1. Plant materials and field experiments

The study was conducted over four consecutive years (2019, 2020, 2021 and 2022), in twelve locations spread across Austria, France, Romania, and Spain. All locations selected for model development and validation needed to be easily accessible and legally eligible for drone use. The main dataset used to define and refine the model was taken in 2020 and 2021. In 2020, data was taken in one location in Austria, one location in France, and one location in Spain. We discarded two locations: one in Spain due to bract necrosis infection and one in France due to experimental design requirements. The 2021 data was taken in one location in Austria, two locations in Spain, and two locations in Romania. All locations were included in the analysis. Additional data from 2019, 2020, and 2022 was included to validate the model. The additional 2019 data consists of one location originally flown as a feasibility study. It was not used for model development due to data integrity issues that were later resolved. The 2020 trial excluded due to design constraints, was reinstated for validation. The added 2022 dataset consists of one location in France, one in Romania, and one in Spain. The total breeding plots evaluated each year according to the GT data (FLW50_GT) and DL methods deployed to estimate 50% of flowering (FLW50_UAV_plcnt and FLW50_UAV_maxfl) are represented in Table 1.

**Table 1.** Number of observations in each trial for the variables: 50% flowering using GT data (FLW50_GT), remote sensing (RS) data modeled using previously captured plant number per plot (FLW50_UAV_plcnt), and remote sensing (RS) data modeled using max number of sunflower heads per plot detected during the flowering flights (FLW50_UAV_maxfl).

| Trials | FLW50_GT |  |  |  | FLW50_UAV_plcnt |  |  |  | FLW50_UAV_maxfl |  |
| --- | --- | --- | --- | --- | --- | --- | --- | --- | --- | --- |
|  | 2019 | 2020 | 2021 | 2022 | 2019 | 2020 | 2021 | 2022 | 2020 | 2021 |
| 1 | - | 1684 | - | - | - | 1677 | - | - | - | - |
| 2 | - | 96 | - | - | - | 96 | - | - | - | - |
| 3 | - | 1897 | - | - | - | 1888 | - | - | 1894 | - |
| 4 | - | - | 1645 | - | - | - | 1591 | - | - | 1591 |
| 5 | - | - | 700 | - | - | - | 568 | - | - | 568 |
| 6 | - | - | 1227 | - | - | - | 1226 | - | - | 1227 |
| 7 | - | - | 2061 | - | - | - | 2061 | - | - | - |
| 8 | - | - | 147 | - | - | - | 147 | - | - | 147 |
| 9 | - | - | - | 39 | - | - | - | 39 | - | - |
| 10 | - | - | - | 704 | - | - | - | 704 | - | - |
| 11 | - | - | - | 765 | - | - | - | 765 | - | - |
| 12 | 1020 | - | - | - | 1020 | - | - | - | - | - |
| **Grand Total** | 1020 | 3677 | 5780 | 1508 | 1020 | 3661 | 5593 | 1508 | 1894 | 3533 |

The climate in the regions where the trials were conducted can be described according to the Köppen climate classification as either Cfb (France, Austria; Temperate, no dry season, warm summer), Dfb (Romania; Cold, no dry season, warm summer) or Csa, (Spain; Temperate, dry summer, hot summer) (Geiger 1954 and 1961). No trials were irrigated or sprayed with post-emergence herbicides. Most fields were protected from animal damage by using fencing and bird repellant techniques. Double plantings, where two seeds are planted very close together, are manually removed to ensure appropriate spacing between plants.

The sunflower germplasm in the selected trials represented genetically diverse hybrids at various stages of the product development process, which were adapted to the local management environment. The experimental design used in the trials was an augmented experimental design with diagonal checks placed throughout the location to provide replication for residual error estimation. There were repeated entries within each trial, across years, and across locations.

### 2.2. Unmanned Aerial Vehicle (UAV) Image Acquisition

UAV-based image acquisition aligned with pivotal growth stages of sunflower crops to ensure comprehensive documentation of the entire flowering period. The UAV was outfitted with a high-resolution RGB camera and flight parameters (altitude, flight speed, photo overlap) were tailored to achieve high image quality data capture to facilitate downstream analysis (Bongomin et al., 2024; Kim & Sung, 2024; Mesas-Carrascosa et al., 2016).

Flights were scheduled around noon local time to mitigate the potential for shadows and variable lighting which could affect image consistency and analysis. An early season flight was conducted to establish the RS plant count. The UAV flights ranged from 3 days before early checks until all plots reached 50% flowering, thereby capturing the full spectrum of the flowering cycle. At least five flights were conducted at each location to document the entire flowering period. The scheduling of flights, from pre-peak to post-peak flowering, provided the high temporal resolution necessary for detailed phenological analysis of sunflower development. The optimal number of flights in this study was optimized based on a previous analysis using a dataset from 2019 where seven flights were flown. The optimum flight number and cadence that had the most value, i.e. a good prediction fit without excess workload, was chosen.

### 2.3. Ground Truth Data Collection, Time Series Analysis and Flowering Date Estimation

Ground-truth (GT) data were collected by human evaluators concurrent with UAV image acquisition. Observers recorded the date when 50% of the plot was flowering by counting the number of plants with heads that have reached the R-5.1 sunflower growth stage. The R-5.1 growth stage is the beginning of flowering in which 10% of the head area has completed or is in flowering (Schneiter & Miller, 1981). These observations provided a basis for calibrating the UAV-based image analysis and for validating the model’s accuracy. Flower counts obtained from the UAV imagery and ground observations were compiled into a time series dataset. The dataset was then analyzed using a logistic regression model to estimate the 50% flowering date (FLW50_GT). The logistic model allowed for the assessment of the probability of a plant reaching the flowering stage at different time points throughout the season (Lever et al., 2016; Maalouf, 2011; Yoosefzadeh Najafabadi et al., 2023).

We employed a logistic regression analysis using the percentage of sunflower flowering heads observed in each plot to synthesize the time-series imaging data and the phenotypic predictions obtained from the CNN. This statistical approach modeled the progression of flowering over time so we could pinpoint the precise date the logistic curve intersected the 50% flowering threshold. This intersection date was designated as the flowering date for each plot, consistent with the traditional agronomic definition where 50% of the sunflower heads have reached the R-5.1 development stage. The estimated flowering time is the intersection between the line associated with half of the ultimate number of sunflower heads counted and the flowering curve. We used this method to translate continuous phenological observations into a discrete, actionable data point for the peak flowering stage for sunflowers.

### 2.4. Sigmoid curve to predict sunflower flower and flight frequency

We employed a nonlinear logistic model to accurately estimate the time of 50% flowering in sunflowers. This choice was motivated by the logistic function’s efficacy in modeling biological growth processes, capturing the sigmoidal pattern of sunflower development in relation to day of the year (DOY). We estimated the number of plants via RS (FLW50_UAV_plcnt) by counting plants at early growth stage and/or estimating the max number of sunflower heads detected (FLW50_UAV_maxfl) by the CNN-DL model.

The logistic model, characterized by its S-shaped curve, can be defined as:

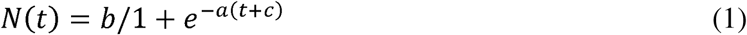

where ‘*N(t)*’ is the estimated number of flowers at time *t* (DOY), ‘*a*’ represents the growth rate, which indicates how rapidly the flowering progress occurs, ‘*b*’ is either the stand count determined by a early flight or the maximum number of flowers observed which serves as an upper limit in the model, ‘*c*’ shifts the curve along the DOY axis, effectively determining the DOY at which 50% of the plants are expected to bloom, and *t* denotes the DOY. The values of ‘*a*’, ‘*b*’, and ‘*c*’ are optimized so that they best fit the observed data using statistical techniques. The model iteratively adjusts these parameters then evaluates the goodness of fit by calculating the overall difference between the observed data points and the model’s predictions across all DOYs considered in the study.

A simulation strategy was implemented to assess the optimal UAV flight frequency for the analyzed data set. Optimal flight frequency was defined as one that would allow accurate data collection and elevate operational efficiency. This involved evaluating a range of random flight datasets subsets, each having three to seven flights, coming from the 2019 feasibility trial. Each dataset subset was analyzed for correlation between GT data (FLW50_GT) and UAV-derived estimates (FLW50_UAV) and goodness-of-fit to the model. Additionally, to refine the FLW50 predictions using the logistic regression model, a threshold was established according to the maturity estimated for late materials to exclude any predictions exceeding 100 days. This cutoff was implemented to address the typical issue of overestimation observed with sparse time-serie data, ensuring more reliable and accurate modeling outcomes.

Preliminary analysis suggested that conducting five flights achieves a high level of accuracy, and a well-fitted logistic curve, as evidenced by a correlation coefficient (*r*) exceeding 0.92, and low estimation errors with a mean absolute error (*MAE*) below 1.16 (Figures 1-a and 1-b). These findings indicate that five flights, scheduled at intervals of around 4 days, yield optimal results comparable to those obtained from seven flights. However, the optimal frequency of flights may vary depending on the nature of the trial. Based on the present results and the available data set, this conclusion appears to be ideal, but different maturity groups or trial weather condition could achieve better performance by selecting a different number and frequency of flights. Furthermore, it is posited that four strategically timed flights, if scheduled to adequately capture the progressive stages of the S-shaped growth curve, have the potential to produce satisfactory outcomes (Supplementary Figures 1 and 2).

**Figure 1-a.**
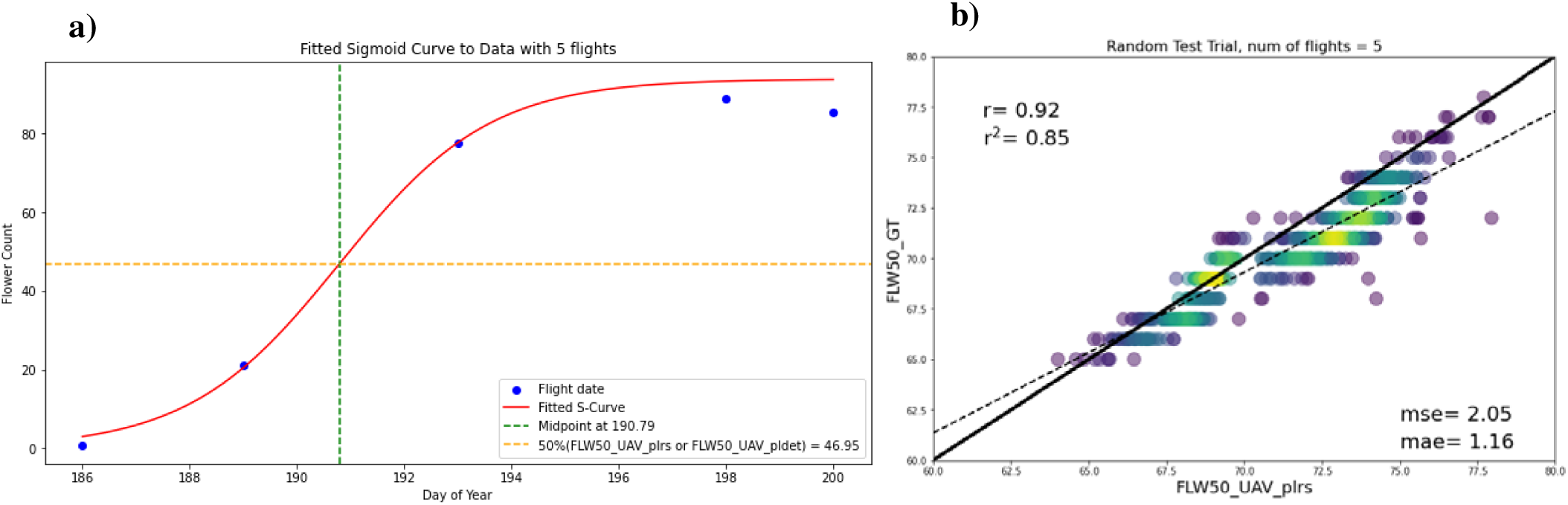
displays the logistic curve fitted using data from five UAV flights, while Figure 1-b assesses the model’s performance relative to this identified optimal flight frequency.

### 2.5. Image Processing and Deep Learning Model for Flower Counting Analysis

The raw images captured by the UAV throughout time were preprocessed to generate georeferenced orthomosaic from each individual flight. A deep learning model was developed to detect and estimate the number of sunflowers in images using density-based approach, commonly applied in crowd counting. To train models using this approach, images needed to be manually annotated with dots marking the sunflower heads. These dot annotations wer convolved with a normalized Gaussian kernel to generate density maps, so that integrating over the density map returns the count of flowers (annotated dots, Figure 2).

**Figure 2:**
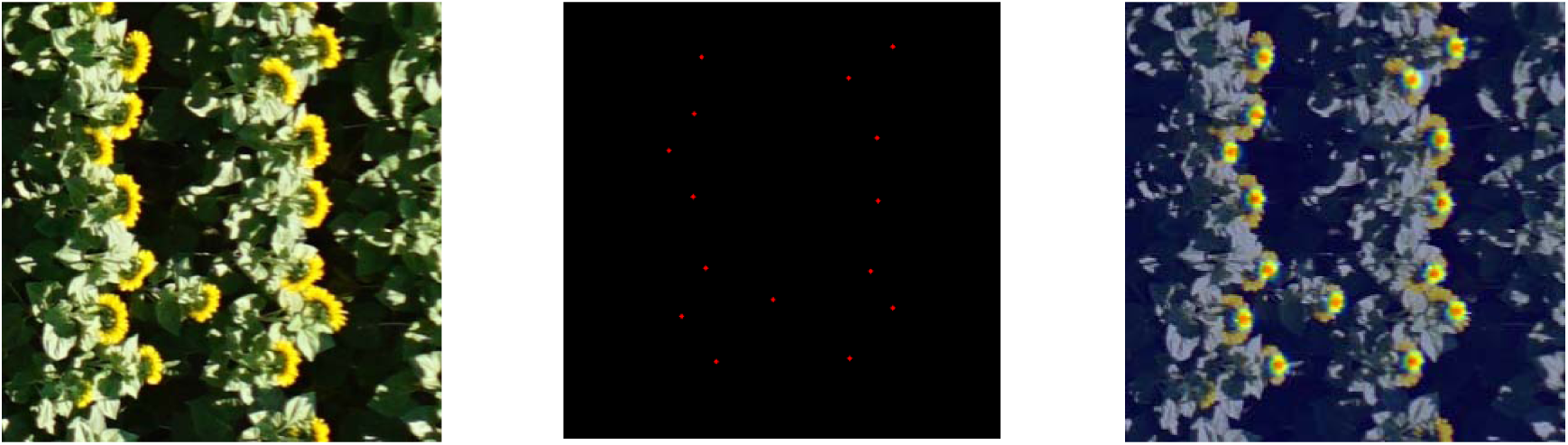
An example image patch (left), its annotation (middle) and visualization of the density map (right).

The model architecture employed was a U-net (Ronneberger et al., 2015), a Fully Convolutional Network commonly used for image segmentation. U-Net is a deep learning network with an encoder–decoder architecture. The encoder uses a series of convolution and pooling layers to down sample and encode the input images into multiple feature representations while the decoder uses up-sampling and convolution layers to transform these features back into high-resolution output, in this case, the density maps. Additionally, there are skip connections that concatenate layers from encoder with layers in decoder to pass on information learned in encoder layers to decode layers.

The dataset consisted of 4,366 image patches and corresponding density maps, each 416 x 416 pixels in size, collected from 10 distinct locations in 2020. This dataset was divided into training and validation subsets following an 80:20 ratio to train for U-net model parameters. During inference, the trained model processed new images to generate density maps. These density maps were then integrated (summing up values) to predict count of sunflowers.

### 2.6. Validation of the Methodology and Statistical Analyses

Validation analysis using the 2020 and 2021 datasets was conducted to ascertain the precision of the model in estimating 50% flowering dates through RS derived plant count (FLW50_UAV_plcnt) and the maximal detection of sunflower heads (FLW50_UAV_maxfl). This was achieved by performing statistical comparisons between UAV-based flowering date estimates and conventional GT measurements. The Pairwise Games-Howell test was used to compare the group means between the observed data, utilizing the default parameters from the R package ‘ggstatsplot’ (Patil, 2021). Discrepancies and congruence were examined through graphical boxplot visualization, correlation coefficients and error metrics to identify both the strength of the relationship and the presence of any systematic biases. Specifically, Pearson’s correlation coefficient (*r*) was utilized to evaluate the linear association between the UAV and ground-measured data. Additionally, the model’s accuracy to estimate flowering date was quantified using the coefficient of determination (*R²*), *MAE*, mean squared error (*MSE*) and Root Mean Squared Error (*RMSE*) providing a comprehensive assessment of the model’s performance and its applicability in practical RS applications. UAV-derived results were also compared with human-derived GT data, using box plots, histograms, and Bland-Altman plots to evaluate data distributions. This approach provided a comprehensive visual analysis, highlighting variations and consistencies across different trials, and effectively illustrating the accuracy and reliability of UAV technology in capturing phenological data.

The most effective method for estimating FLW50 using UAV-derived data was then selected to advance the analysis, incorporating trials from 2019 and 2022. To optimize model performance for prediction the means for the evaluated trials a mixed model was deployed to generate BLUPs (Best Linear Unbiased Predictions) and BLUEs (Best Linear Unbiased Estimates), as well as genotypic components with spatial structures. Single-environment trial models, adjusted using the package ASReml-R 4 (Butler et al 2017), were fitted for each genotype in each environment as follows:

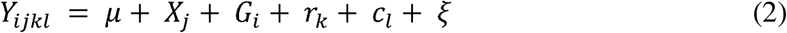

where *Y_ijkl_* is the observed 50% to flowering date of the *i^th^* genotype of the *j^th^* experiment with diagonal checks, *k^th^* row and *l^th^* column; *μ* is the fixed vector represented as 1 for the general mean; *X_j_* is the fixed effect of the *j^th^* experiment; *r_k_* is the random effect of the *k^th^* row, with 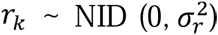, where 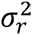 is the variance of rows effects; *c_l_* is the random effect of the *l^th^* column, with 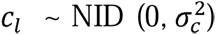, where 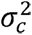 is the variance of columns effects; *ξ* is a vector of spatially dependent or correlated random errors in the row-column structure, where, *ξ* ∼ N (0, R) and 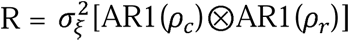. R represents the Kronecker product between separable auto-regressive processes of first order in the row-column dimension; 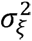 is the variance of spatially dependent residue or due to the spatial trend; and *ρ_c_* and *ρ_r_* are the autocorrelation parameters for column and row, respectively (Cullis et al., 2006). The predicted genotype means are obtained by solving the mixed model equation. The random effects of the models were tested by likelihood ratio test (Rao, 2002) considering a chi-square distribution with one degree of freedom

The model’s efficacy was further evaluated through the calculation of the individual broad-sense heritability 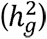 according to Cullis et al. (2006), by the formula:

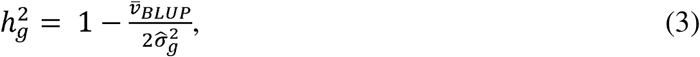

where 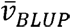 is the average variance of pairwise differences between the BLUP of genotype effects. Additionally, the accuracy 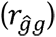 was also estimated as following (Hickey et al., 2009):

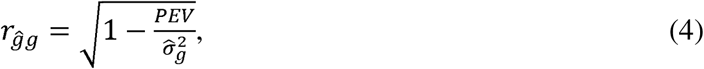

where *PEV* is the prediction error variance, obtained by the diagonal elements of the inverse of the left-hand side of mixed models equation. Finally, the performance assessment was enriched by analyzing genotypic components in each trial, providing a comprehensive view of the model’s ability to capture genetic influences on estimate flowering dates using UAV techniques.

## 3. RESULTS

### 3.1. Accuracy of validation method

The influence of different environments on flowering date estimations can be highlighted by the range of flowering times observed between trials. The flowering date across trials for GT measurements (FLW50_GT) and UAV-based estimations ranged from 66 to 94 days after planting (DAP) and a similar data distribution was observed in all environments (Figure 3). The overall mean by trial showed that trial three presented the longest flowering dates for all flowering date methods, exceeding 84 DAP. Conversely, the shortest flowering dates were observed in trial six falling below 76 DAP.

**Figure 3:**
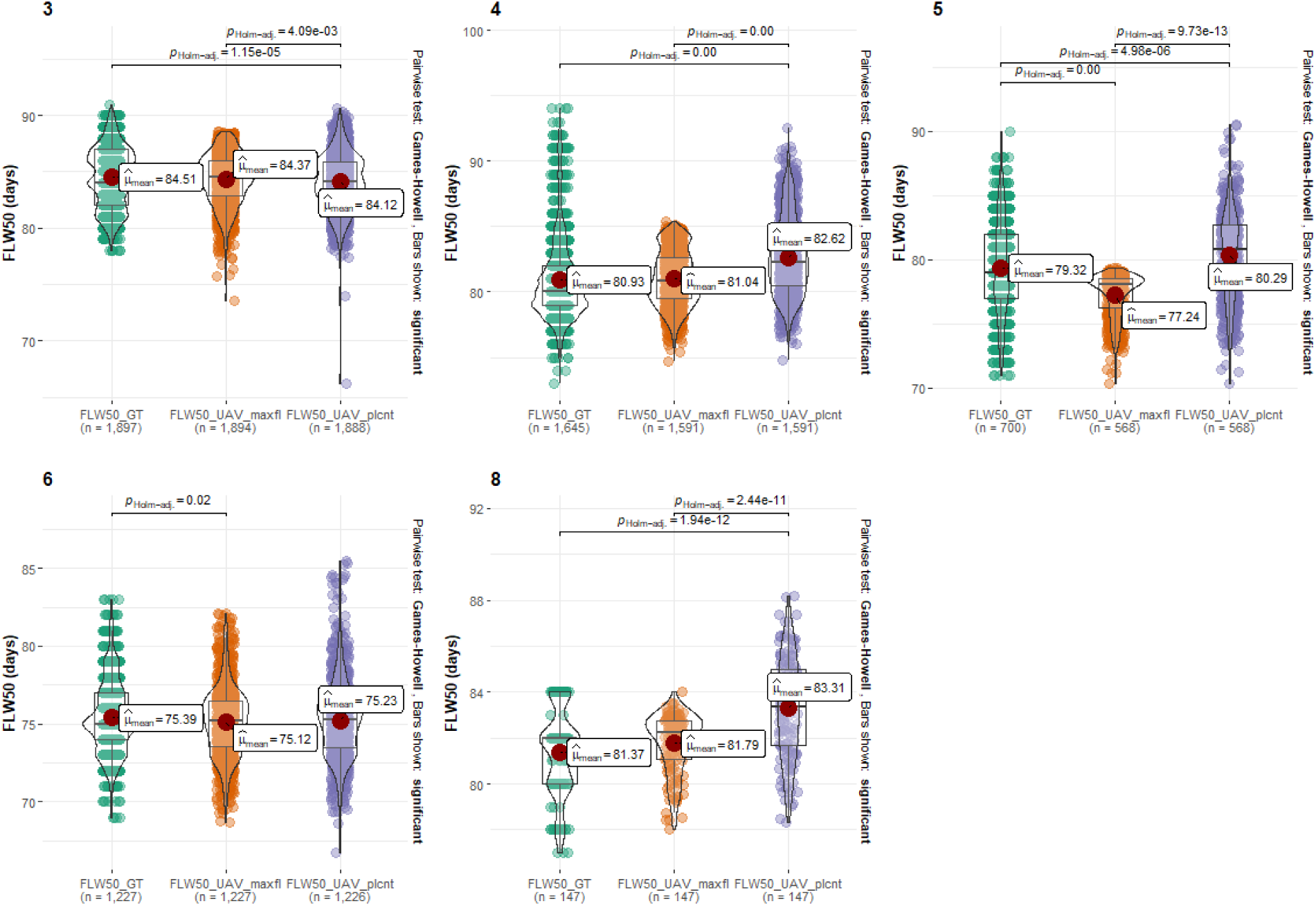
Combination of box and violin plots along with jittered data points for between subjects comparisons by trials to flowering date (FLW50) from human-derived GT and UAV-derived data using maximum number of detected sunflower heads (FLW50_UAV_maxfl) and UAV stand count (FLW50_UAV_plcnt). The data encompasses one trial from 2020 (trial 3) and four trials from 2021 (trials 4, 5, 6, and 8). Group means were compared using the pairwise Games-Howell test. Comparisons show only significant values. The data is segmented by trial number, illustrating the variability and central tendency of each variable’s measurements, as well as the significancy value according to the deployed test.

In the analysis of all validation datasets, the distribution of the FLW50_UAV_plcnt data closely paralleled that of the FLW50_GT in all validation trials, with notable exception in trial eight. However, the Pairwise Games-Howell test used to compare means showed disparities (*ρ_Holm_adj_* < 0.05) between FLW50_UAV_plcnt and FLW50_GT values in all trials except trial six. These findings suggest potential deviations in UAV imagery effectiveness or GT counting accuracy in the specific trials where discrepancies were observed. The FLW50_UAV_maxfl data showed trends similar to the FLW50_GT measurements throughout all trials, though with visibly less variability in validation trials four and five. However, in trial five and six, significant differences were observed between FLW50_UAV_maxfl and FLW50_GT, with the disparity being more pronounced in trial five (Figure 3). This particular condition in trial five can be illustrated by examining the data distribution across different window size ranges for DAP. Supplementary Figure 3 highlights the similarities between the bins by displaying the range size for each deployed method of flowering estimation date. Close alignment between the UAV-derived data and FLW50_GT in most trials, as depicted in Figure 3, indicates overall effectiveness of the UAV approaches, while specific deviations for a particular DAP range are shown in Supplemental Figure 3.

The Bland-Altman plots provided in Supplementary Figure 4, offer a statistical assessment of the agreement between UAV-derived flowering date estimates and GT evaluated data. These plots detail the mean differences and limits of agreement for each trial, showing the consistency of UAV-derived measurements with human-derived field observations. The analysis showed low variability in the predicted UAV flowering dates in each trial with a significance level of 95% confidence intervals (CI), which serves as a critical measure of the precision of UAV estimates. For instance, the CI for FLW50_UAV_maxfl reaches a high of 3.6% in trial 3, indicating a greater variability in estimates, whereas it drops to a low of 0.3% in trial 8, suggesting a tighter agreement with GT. Similarly, FLW50_UAV_plcnt exhibits a maximum CI of 3.4% in trial 4, with a reduction to 1% in trial 8.

Results comparing UAV-derived estimates for the 50% flowering date against human-derived GT observations across various trials demonstrated strong correlations and low error rates, indicating a robust performance of UAV methodologies (Table 2). The correlations of both UAV-derived methods with GT data were consistent. The highest levels of correlation for both UAV-based methods were recorded in trials three and six, with a correlation coefficient (*r*) exceeding 0.95. These trials also indicated low error metrics, registering less than 1. Notably, in trial five, the results of FLW50_UAV_plcnt exhibited superior accuracy compared to those obtained from FLW50_UAV_maxfl, underscoring the potential for method-specific efficacy within certain trial conditions. The analysis of the relationship between FLW50 results and the accuracy of UAV estimations compared to field data is depicted graphically in Supplementary Figures 5 and 6. Supplementary Figure 5 focuses on FLW50_UAV_plcnt, while Supplementary Figure 6 pertains to FLW50_UAV_maxfl, providing a visual representation that demonstrates how closely UAV-derived estimations align with actual field measurements.

**Table 2:**
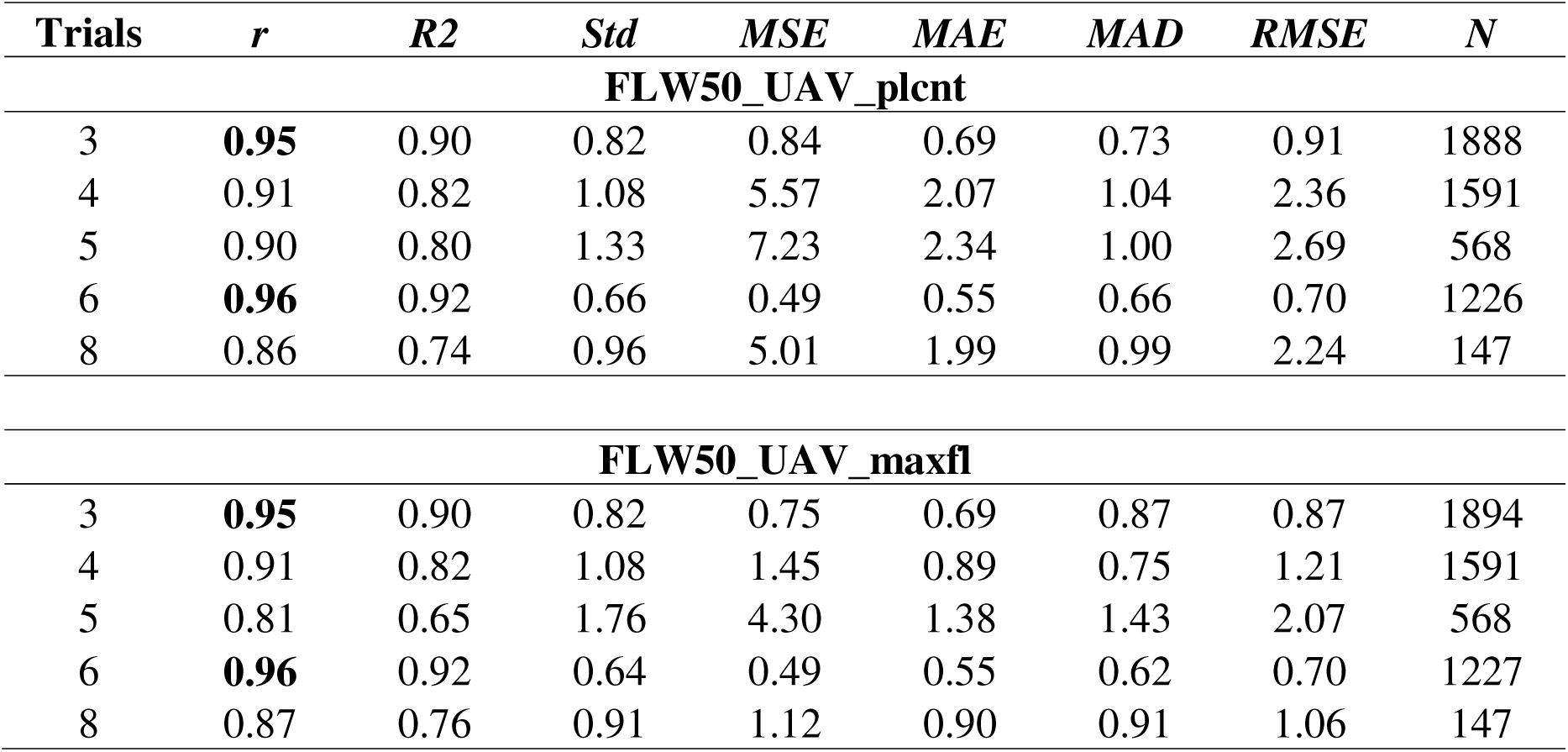
Statistical Analysis of Flowering Date Estimations from UAV-Derived Data. This table summarizes the correlation coefficient (*r*), coefficient of determination (*R²*), standard deviation (*Std*), mean squared error (*MSE*), mean absolute error (*MAE*), mean absolute deviation (*MAD*), root mean squared error (*RMSE*), and the sample size (*N*) for UAV-derived plant stand counts (FLW50_UAV_plcnt) and max head flower detections (FLW50_UAV_maxfl) by trials.

### 3.2. Model performance to flowering dates

Incorporating 2019 and 2022 datasets, the observed FLW50 across trials exhibited a range from 55 to 94 ± 0.06 DAP for FLW50_GT, while the range was slightly broader for FLW50_UAV_plcnt, spanning from 58 to 92 ± 0.06 DAP. Trial 11 presented the earliest mean flowering dates, while the latest mean flowering dates for both variables was in trial three claimed. The boxplots in Supplementary Figure 7 provide a visual summary of the distribution of the 50% flowering dates across the various trials using the complete data set (validation and inference) from all years.

We tested the model’s performance with GT data by using FLW50_UAV_plcnt to estimate flowering dates based on the validation results. The scatter plot analysis showed strong correlations with GT measurements across all trials, indicating that the UAV methodology provides reliable estimates of 50% flowering dates (Figure 4). Trials three, six, and eleven demonstrated particularly robust correlations, with correlation coefficients (*r*) exceeding 0.95, suggesting a close relationship between the UAV data and GT observations. The lowest correlation in our study was observed in trial 10, where the correlation coefficient was 0.84. This trial also exhibited multiple outlier data points, which likely contributed to the reduced correlation observed. Furthermore, error metrics like mean absolute error (*MAE*) and root mean square error (*RMSE*) were generally low, further demonstrating UAV-based estimation accuracy. These findings underscore the effectiveness of using UAV technology for phenological tracking of flowering in sunflower cultivation, providing a valuable tool for precision agriculture and plant breeding programs.

**Figure 4:**
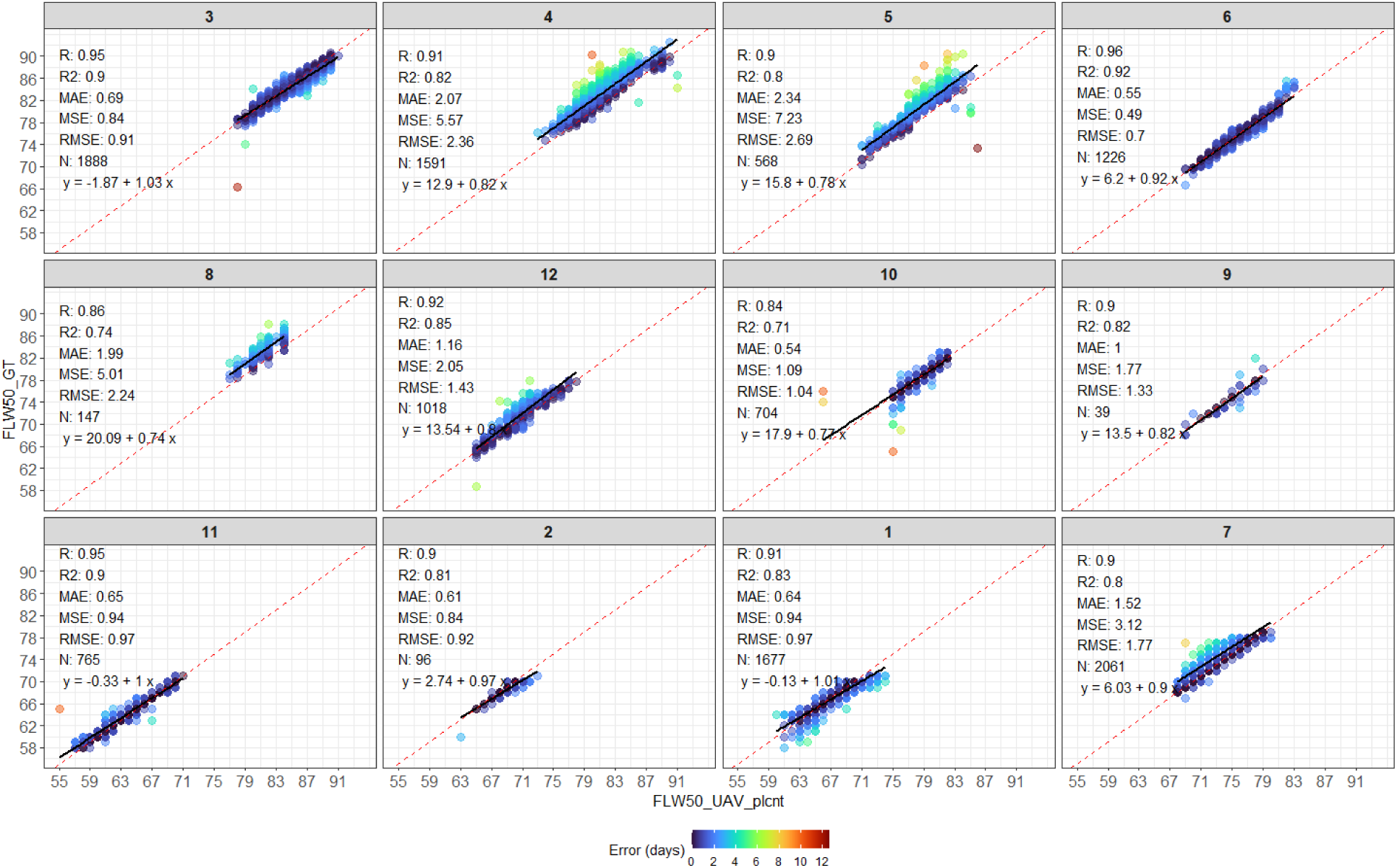
Correlation and error analysis of 50% flowering date estimations for sunflower trials. The scatter plots illustrate the relationship between the ground truth 50% flowering date (FLW50_GT) and UAV-derived estimates using plant stand counts (FLW50_UAV_plcnt) across 12 different trials. The color gradient indicates error magnitude in days. Statistical metrics such as correlation coefficient (*R*), coefficient of determination (*R²*), mean absolute error (*MAE*), mean squared error (*MSE*), root mean squared error (*RMSE*), and linear regression equations are provided for each trial, demonstrating the accuracy of UAV estimations against field data.

### 3.3. Flowering date estimation and accuracy of genetic components

The genetic component’s influence on the 50FLW relative to GT and UAV-derived measurements exhibited a statistically significant impact across the majority of trials (Supplementary Table 1). However, trial 9 deviated from this trend, not showing a significant genotypic effect across both variables. This anomaly can be attributed to the low number of observations and the unbalanced dataset used in the mixed model analysis. Similarly, data from trial 8 also did not present a significant genetic effect when analyzed using the full mixed model (Eq. 2, with *P*-values > 0.01). However, an alternative mixed model approach without spatial correction structure was utilized for this specific trial to estimate the components of variance and predict the genotypic values.

The analysis of 50% flowering dates across trials, as illustrated by the BLUEs in Figure 5, highlights the range of phenological responses and resembles the distribution found in the raw data set. Notably, trial 11 exhibited the earliest average flowering dates, while trial 3 showed the latest. When evaluating the consistency of traditional ground measurement methods against UAV-based estimations, the results were congruent across the board. The correlation coefficients for BLUEs values by trials were notably strong, with all correlations exceeding an *r*-value of 0.81, demonstrating consistent agreement between measurements and estimations for a given trial as detailed in Supplemental Figure 8. This congruence suggests that UAV-derived data can reliably mirror phenological trends, offering a valid alternative to traditional ground measurements for monitoring flowering patterns across diverse maturity groups of sunflower varieties.

**Figure 5:**
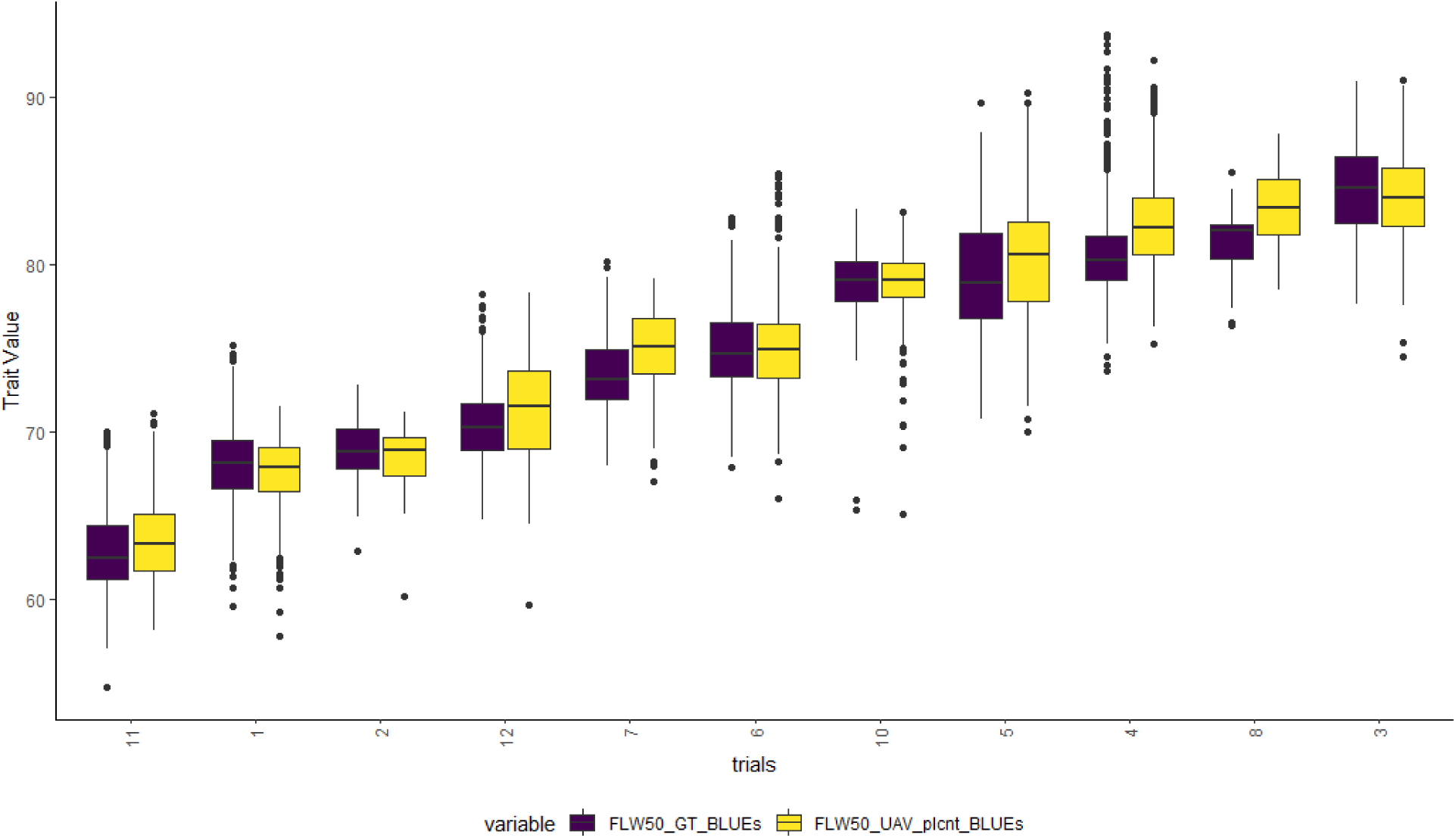
Comparative BLUEs-Based Distribution of 50% Flowering Dates Across Trials. The boxplots depict both the GT dates (FLW50_GT) and UAV-derived plant stand count estimates (FLW50_UAV_plcnt), highlighting the data’s spread and central tendencies for each trial conducted within the study.

Genetic effects 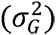 were the predominant contributors to the variability observed in both GT flowering dates (FLW50_GT) and UAV-derived stand counts (FLW50_UAV_plcnt), accounting for over 63% of the variance in all trials. The comparative analysis of genetic parameters between these two variables revealed a consistent alignment or even an enhancement when using UAV-derived data, with trial five being the sole exception. Similar patterns can observe to heritability parameters 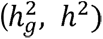, accuracy 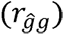 and variance error 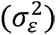 underscoring the UAV-derived outputs in the predictions of sunflower dates in all trials evaluated in this study (Figure 6). In general, heritability is consistent across trials, with occasional variations in error rates and genetic variance, which may reflect trial-specific environmental influences.

**Figure 6:**
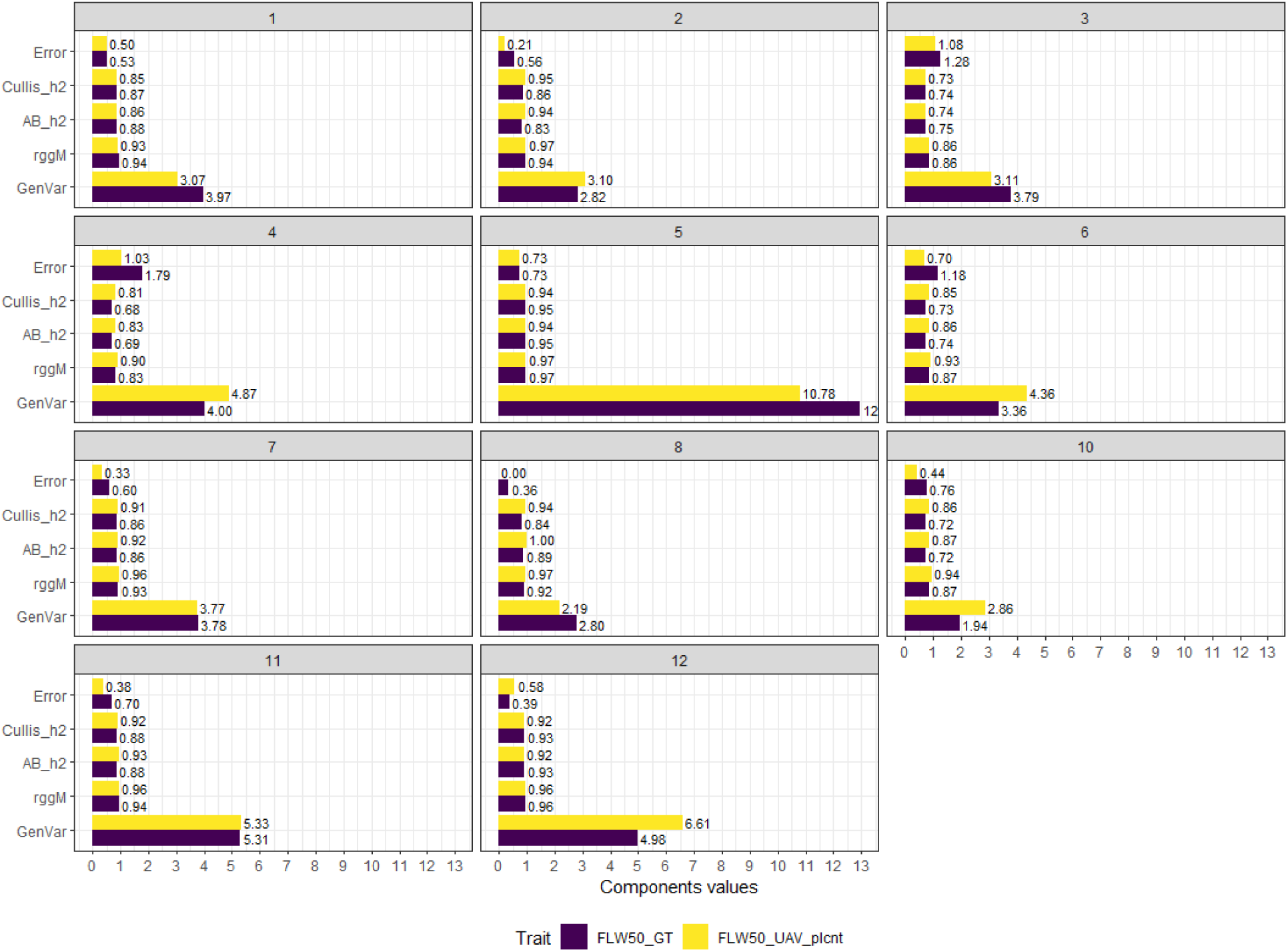
Heritability and Genetic Variance Component Analysis Across Trials. Each panel illustrates the values for error rate 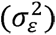, Cullis heritability 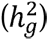, animal breeding heritability (*h*^2^), genetic accuracy 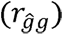, and genetic variance 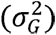 associated with GT (FLW50_GT) and UAV-derived 50% flowering date estimates (FLW50_UAV_plcnt).

## 4. DISCUSSION

### 4.1. UAV Time series data in plant breeding

UAV platforms can facilitate assessment of traits over time, capturing the dynamics of plant development. This temporal insight can be crucial in phenology, especially in assessing flowering times, which are crucial to understanding the interaction between genetics and environment. Numerous studies have highlighted the pivotal role that temporal dataset, acquired through RS methods, plays in modern agriculture for the assessment of breeding traits (Diao & Li, 2022; Sishodia et al., 2020; Turkoglu et al., 2021). By elucidating the genetic basis of environmental adaptability in sunflower crops during flowering growth stage, UAV-derived data can guide the selection of traits that optimize plant performance under diverse conditions. This is especially pertinent in devising sunflower agricultural practices that align with environmental constraints, such as sowing date and plant density (Barros et al., 2004), optimizing irrigation schedules (Göksoy et al., 2004) or determining the appropriate timing for applying growth regulators (Spitzer et al., 2018) (SPITZER et al., 2018). HTP methods are gaining traction in agricultural research but their application in sunflower breeding, especially involving analyses at multiple time points, has been less extensively documented. Sunflower phenological traits, such as flowering timing and pattern, could benefit from understanding the temporal insights provided by RS approaches.

UAVs enable frequent and consistent monitoring, which is critical for capturing the temporal variability inherent in crop development and for identifying traits that confer advantages under specific environmental conditions. In this study, we demonstrated that the flowering date could be estimated with high accuracy across several environments (*r* > 0.84), compared to traditional field assessments. Moreover, this method yielded even better results when analyzing heritability values in several trials (Figure 6). In studies involving corn, the use of time-series UAV-based hyperspectral data demonstrated a correlation of 0.92 between the UAV estimates and the actual GT for the days to anthesis using ridge regression machine learning model (Fan et al., 2022). However, the findings were derived from analyzing full-band reflectance patterns rather than employing a counting method to determine the actual number of plants reaching the flowering stage as performed in the present study. Another study was conducted to estimate flowering time in sorghum fitting a third-degree polynomial function using the estimated counted panicle (Cai et al., 2021). As in our study, the authors found good estimation precision and low error, but a limited data set was analyzed which made it difficult to generalize the model to other environments in sorghum fields.

The investigation of UAV-time-series data to measure physiological traits such as plant maturity and plant health, nitrogen efficiency, and others also have been largely investigated using UAV-imagery data in crops (Feng et al., 2021). In soybean, date of maturity is an important trait that successfully has been validated using UAV time-series data not only using images from drone (Moeinizade et al., 2022; Trevisan et al., 2020; Volpato et al., 2021) but also satellite (Mazis et al., 2023). The correlations for soybean breeding plots between GT measurements and maturity estimates in these studies ranged from 0.6 to 0.9 with an average ± 2 days off. Our results indicated improved outcomes when considering plant maturity in soybeans. It is important to note that in this study, we applied a direct DOY estimation by counting the flowering heads using CNNs and DL approaches rather than selecting a threshold to infer a physiological growth stage denoted as maturity through linear regression models.

While CNNs have not traditionally been used for estimating temporal growth dates, recent studies highlight their potential advantages over other methods to predict breeding traits (Teodoro et al., 2021; Xu et al., 2023; Ye et al., 2023). We demonstrated that CNNs can effectively perform sunflower head detections and count during the flowering growth stage as shown by the similar performance between FLW50_UAV_plcnt and FLW50_UAV_maxfl. We were able to achieve enhanced results showcasing the efficiency of CNNs in agricultural applications by integrating CNN-derived data on sunflower head detections with logistic functions. Trevisan et al. (2020) utilized UAV imagery alongside CNNs to predict the maturity date of soybean lines, demonstrating the utility of UAV-based phenotyping systems for commercial breeding programs. The authors validated the proposed method using extensive GT observation from multiple trials which showed a good generalization capability with a RMSE lower than two days in most cases. Moreover, other studies applied a deep learning model combining CNNs and Long Short-Term Memory (LSTM) networks to estimate days to maturity for soybean (Moeinizade et al., 2022) and dry beans (Volpato et al., 2023). The CNN-LSTM was used to determine the accuracy of maturity prediction under varying UAV flight frequencies and environment conditions. Therefore, employing UAV time-series data and CNNs approaches for temporal data assessments is not only feasible but offers significant improvements over traditional methods when measuring phenological traits.

Deep neural networks provide promising results for detecting and counting sunflower heads. Conversely, in cereal crops, DL models have been effectively applied for detecting panicles (Ghosal et al., 2019; H. Li et al., 2022), spikes (Z. Li et al., 2020; Zhang et al., 2022), and tassels (Alzadjali et al., 2021; Jia et al., 2024; Zou et al., 2020), highlighting their potential in agricultural research. The reliability and performance of detecting and counting plant structures using DL models are influenced by several key factors. These include the quality and resolution of the imagery data (C. Hu et al., 2021), variability in plant morphology, the presence of occluding elements such as overlapping leaves or shadows (Kattenborn et al., 2019), and the specific characteristics of the crop species such as wheat (Y. Song et al., 2023), sorghum (Lin & Guo, 2020), and corn (Casuccio & Kotze, 2022). Additionally, environmental conditions at the time of image capture and the complexity of the background terrain can significantly affect model performance in agricultural research (Lu et al., 2024). The DL model utilized in this research for sunflower detection demonstrated robust performance across a variety of environmental conditions, ensuring reliable outcomes. Therefore, plant breeding programs can leverage the reliable outcomes provided by the DL model for sunflower detection to make informed decisions about selecting and breeding sunflower varieties.

### 4.2. Days to sunflower flower accuracy

To ensure accurate detection of sunflower heads using the proposed method, training must be done using high-quality RS images acquired from the field, along with precise human-derived GT data. Compared to field observations, drone imagery must provide good quality images from the target to be able to detect plant features (Hou et al., 2021). In the present study, discrepancies, such as overestimations and underestimations, can be attributed to inaccurate GT data and suboptimal image quality, respectively. A comprehensive investigation using time-series drone imagery to estimate plant maturity in soybeans revealed that almost 50% of outliers stemmed from incorrect field evaluations by human examiners (Volpato et al., 2021). This highlights a need for a similar in-depth analysis in sunflowers to better understand the impact of such factors on the accuracy of aerial imagery assessments.

In addition to image quality and CNN model accuracy, the estimation of sunflower flowering dates may be influenced by factors such as disease, seed dormancy, and planting errors which contribute to the uneven distribution of plants within targeted breeding plots (Dedić et al., 2022; Srivastava & Singh, 2020). Diseases that delay or accelerate flowering due to stress, or deform the flowering head are of particular concern. Instances of severe bract necrosis leads to significant plant stunting, undersized, and scorched flower heads. Detecting bract necrosis poses a particular challenge in aerial imagery analysis due to potential difficulties in visualizing these affected plots from drone-captured images (Figure 7). Occurrence of bract necrosis is difficult to predict as it is not associated with a pathogen (Yang & Berry, 1983) and is generally regarded as a calcium assimilation defect or due to heat and/or moisture stress (Guardia et al., 2008). Downy mildew also deforms the head, but affected plants are fairly detectable in aerial imagery (unpublished data). Even under such adverse conditions, breeding strategies need to account for and evaluate plots. These insights underline the complexity of accurately monitoring sunflower phenology and the importance of integrating comprehensive field observations with UAV imagery analysis.

**Figure 7:**
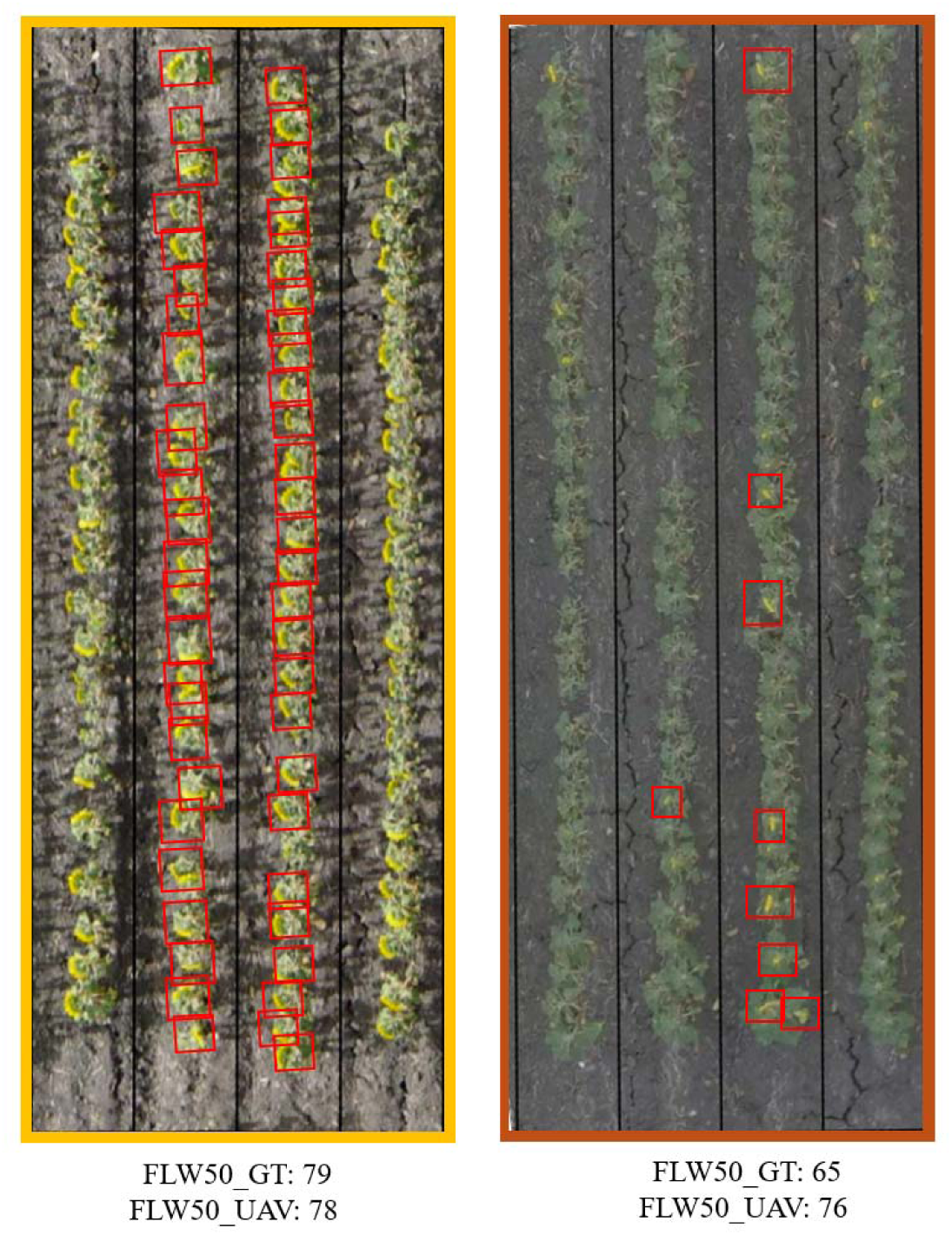
Sunflower breeding plots showing stress in red box (burned heads) and healthy full bloom plants in yellow box. The two central rows were measured in this study. Red boxes represent the detected object (sunflower heads) by the CNN model. This sample is taken from trial 10, referenced in Figure 4.

### 4.3. Logistic model outputs in plant breeding and flight frequency

Logistic models, particularly useful for binary or categorical outcome predictions, offer a better fit for data where the outcome is expected to follow a sigmoid curve, typical in biological processes like flowering, canopy cover and maturity (Hilty et al., 2021; Xavier et al., 2017). Unlike linear regression models that predict a continuous output, logistic models generate probabilities, making them more suitable for cases where the response variable is categorical or when predicting the occurrence of an event within a given timeframe (Nick & Campbell, 2007). Moreover, logistic regression does not require a linear relationship between dependent and independent variables (Chafai et al., 2023). In this study, better results were found when adopted logistic regression rather than linear regression (Table 3) when a comparison analysis was done to a selected trial (referred to trial 12).

**Table 3:** Summary of Regression Model Performance Across Different Flight Numbers. The table displays the coefficients of Pearson correlation (*r*), coefficient of determination (*R2*), mean squared error (*MSE*), mean absolute error (*MAE*), root mean squared error (*RMSE*), and the number of observations (*N*) for linear and logistic regression models.

| Flight number | $r$ | $R^2$ | $MSE$ | $MAE$ | $RMSE$ | $N$ |
| --- | --- | --- | --- | --- | --- | --- |
| **Linear Regression** |  |  |  |  |  |  |
| 3 | 0.64 | 0.40 | 104.58 | 8.21 | 10.23 | 959 |
| 4 | 0.60 | 0.35 | 80.40 | 7.72 | 8.72 | 1005 |
| 5 | 0.73 | 0.53 | 72.55 | 7.20 | 8.52 | 1003 |
| 6 | 0.89 | 0.80 | 148.58 | 9.64 | 12.19 | 824 |
| 7 | 0.88 | 0.78 | 98.70 | 7.88 | 9.93 | 961 |
| **Logistic Regression** |  |  |  |  |  |  |
| 3 | 0.34 | 0.11 | 19.76 | 3.72 | 4.45 | 627 |
| 4 | 0.94 | 0.88 | 1.49 | 0.96 | 1.22 | 994 |
| 5 | 0.92 | 0.85 | 2.05 | 1.16 | 1.43 | 1018 |
| 6 | 0.95 | 0.91 | 1.01 | 0.77 | 1.00 | 1020 |
| 7 | 0.96 | 0.91 | 0.88 | 0.73 | 0.94 | 1020 |

In terms of accuracy, increasing the frequency of UAV flights tends to improve the performance of linear regression models. However, in scenarios using just four flights, logistic regression demonstrated strong correlation with GT data. Comparing both models in the optimal scenario of seven flight time-points revealed that the linear regression model underperformed relative to the logistic model (Figure 8). Furthermore, when error metrics such as *MSE* and *MAE* were evaluated, logistic regression exhibited significant improvements over linear regression across all flight sets used in the leave-out flight’s scenario. The implementation of a threshold to exclude outlier predictions (specifically those predicting flowering days over 100) resulted in the removal of numerous observations in linear regression models at lower flight frequencies, compared to the total number of observations in the test trial (*N* = 1020).

**Figure 8:**
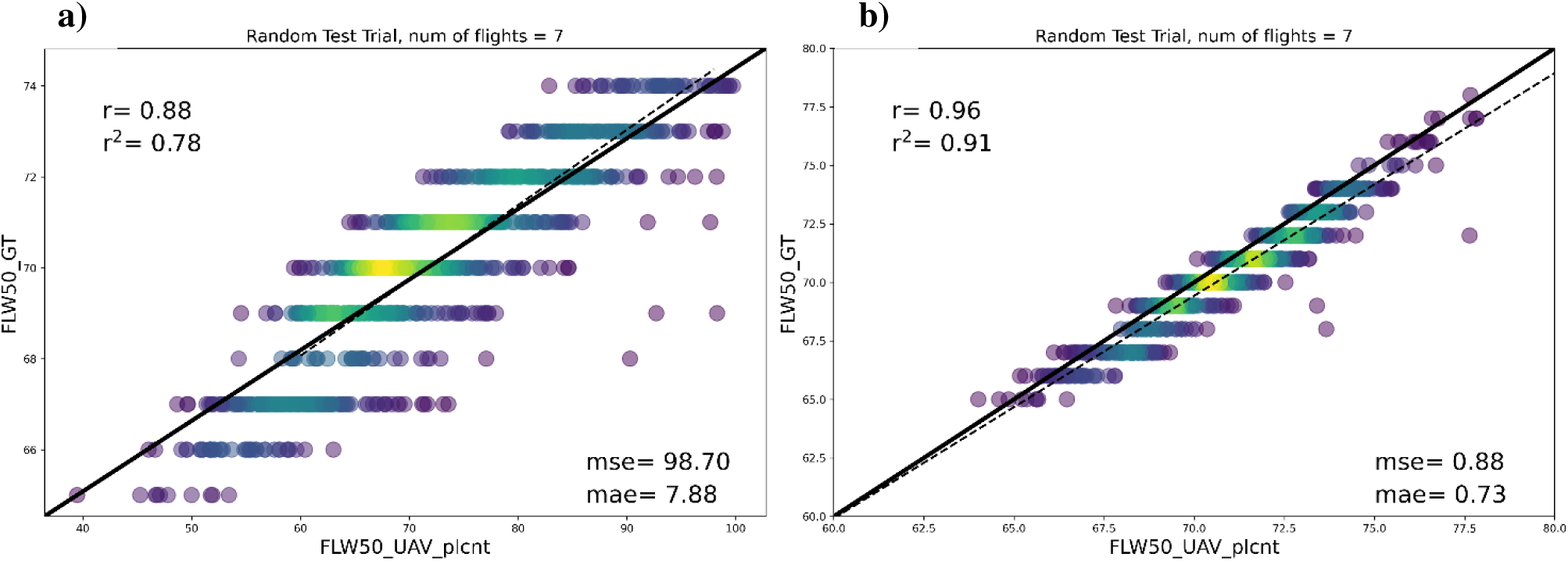
Comparison of Correlation Techniques for 50% Flowering Date Estimation. This figure presents the correlations between GT data (FLW50_GT) and UAV-derived data (FLW50_UAV_plcnt) over seven flights, analyzed using two regression methods: simple linear regression (Panel a) and logistic regression (Panel b). These analyses highlight the effectiveness of each regression technique in predicting the 50% flowering date.

The relationship between flight frequency and model performance will vary depending on the crop growth development and the environment, which includes not only flight conditions but also weather and genetic performance. Increasing the frequency of UAV flights can lead to more accurate and reliable regression models by providing a larger dataset for analysis. This can result in better predictions and insights into crop growth and environmental factors. However, it is essential to carefully consider the specific scenario and type of regression model being used, as the relationship between flight frequency and model performance can vary. In breeding experiments, the window between flights to estimate date of maturity is largely affected by stressed environments, such as dry or excessive heat weather. However, under normal conditions once flight a week has potential to estimate mature with accuracy (Moeinizade et al., 2022; Volpato et al., 2021).

In this study, we have demonstrated that a logistic regression model can effectively estimate the 50% flowering dates in sunflowers using a set of five flights or data points. These data points must capture the plots before any have reached 50% flowering and extend past the date when all plots have reached 50% flowering. For the model utilizing maximum flower counts (FLW50_UAV_maxfl), these flights must continue until all plots have reached 100% flowering, whereas flights can stop earlier if plant counts are used instead. Therefore, it is crucial to maintain a high frequency of data collection throughout the duration of each maturity group to accurately capture the flowering progression.

By using the logistic model outputs from flowering dates estimation, the *"a"* growth rate parameter (Eq. 1) can be correlated to yield in order to investigate genetic relationships between flowering period to grain yield. Early and high yield varieties can be selected by leveraging the growth rate coefficient *"a"* from the logistic model. Short flowering period can affect in many ways grain yield in crops (J. Guo et al., 2021; F. Wang et al., 2023). Higher values of the coefficient *"a"* would typically indicate genotypes with quicker or more intense flowering periods, while lower values might suggest genotypes with more prolonged flowering periods, potentially suitable for selection as late-flowering varieties. By analyzing these relationships, researchers can gain insights into how early flowering traits influence overall crop productivity, facilitating targeted genetic enhancements to improve yield outcomes. In our study, we found no significant relationship between the growth rate and grain yield, as indicated by a correlation coefficient (*r*) of less than 0.05 across most trials. Specifically, in trial 11, where a slightly higher remained low at *r* = 0.15. This suggests that the growth rate may not be a reliable predictor for correlation was anticipated due to early maturity compared to the other trials, the correlation selecting high-yield early genotypes under the conditions tested. Further research could explore alternative variables or factors that could better explain the relationship between flowering period and crop productivity. Furthermore, studying the influence of other environmental factors on growth rate, such as temperature and soil quality, could provide valuable insights for future genetic enhancements and the selection of early high-yield varieties.

## 5. CONCLUSION

We demonstrated the feasibility of measuring the 50% flowering stage by counting sunflower heads from UAV images. We evaluated the performance of two methods to estimate plant populations and found that plants at the early growth stage yielded the most accurate results. We obtained accurate measurements of the 50% flowering dates for sunflowers using UAV time-series data combined with deep learning approaches modeled by logistic regression. Overall accuracy of these estimates is closely aligned with manual counting results, reducing labor costs and increasing flowering dates estimates while simultaneously preventing human error. This information is crucial for plant breeding programs as it allows for precise assessment of the growth and development of sunflower varieties.

## 6. ACKNOWLEDGEMENTS

We extend our heartfelt thanks to everyone at Corteva Agriscience who reviewed the manuscript and provided valuable feedback. Their insightful comments and suggestions improved both the scope of our research and the quality of our manuscript.

## 7. SUPPLEMENTAL MATERIALS

**Supplemental Figure 1:**
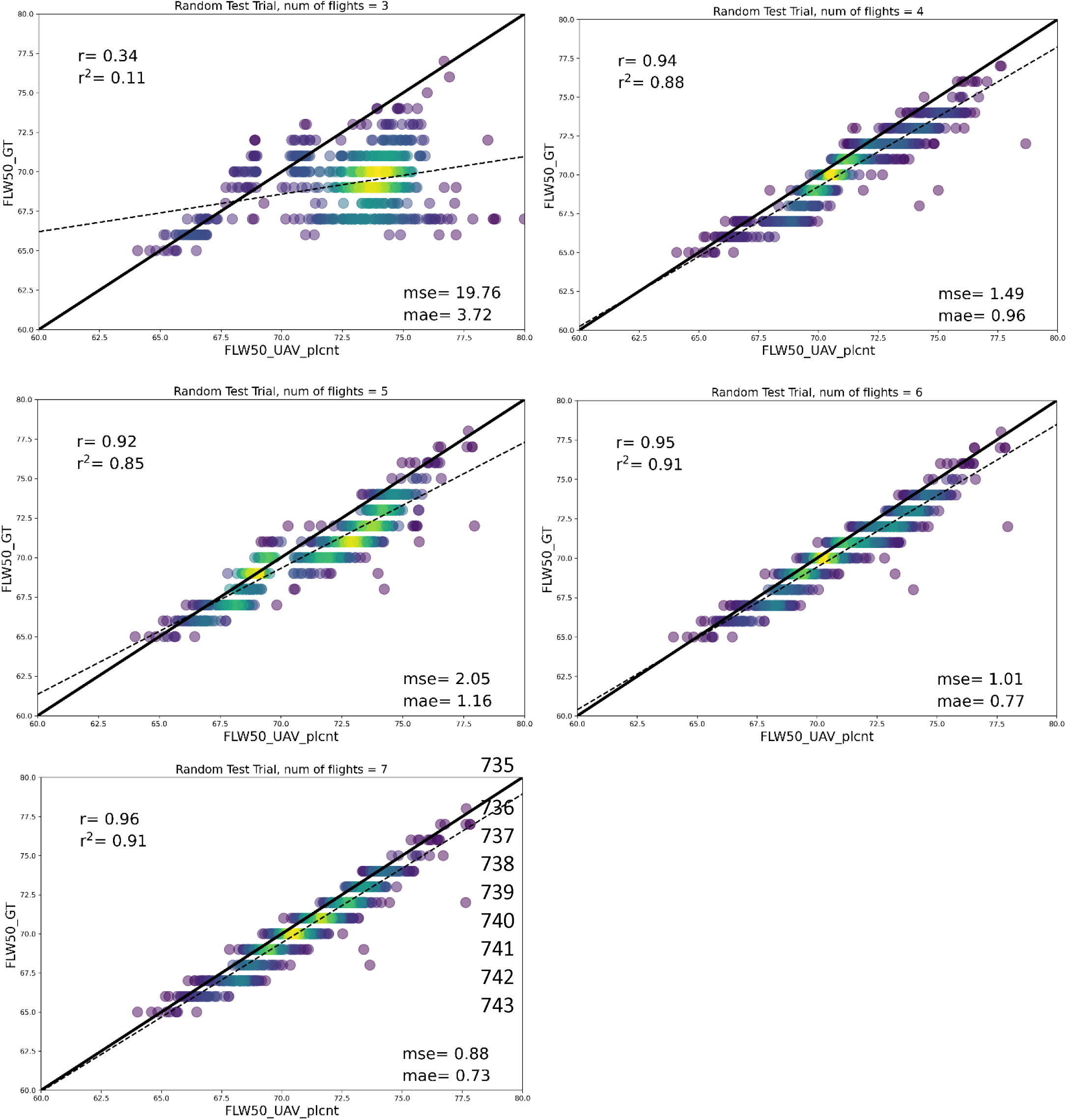
Correlations between GT (FLW50_GT) and UAV (FLW50_UAV_plcnt) 50% flowering date estimation throughout the range of flight frequencies.

**Supplemental Figure 2:**
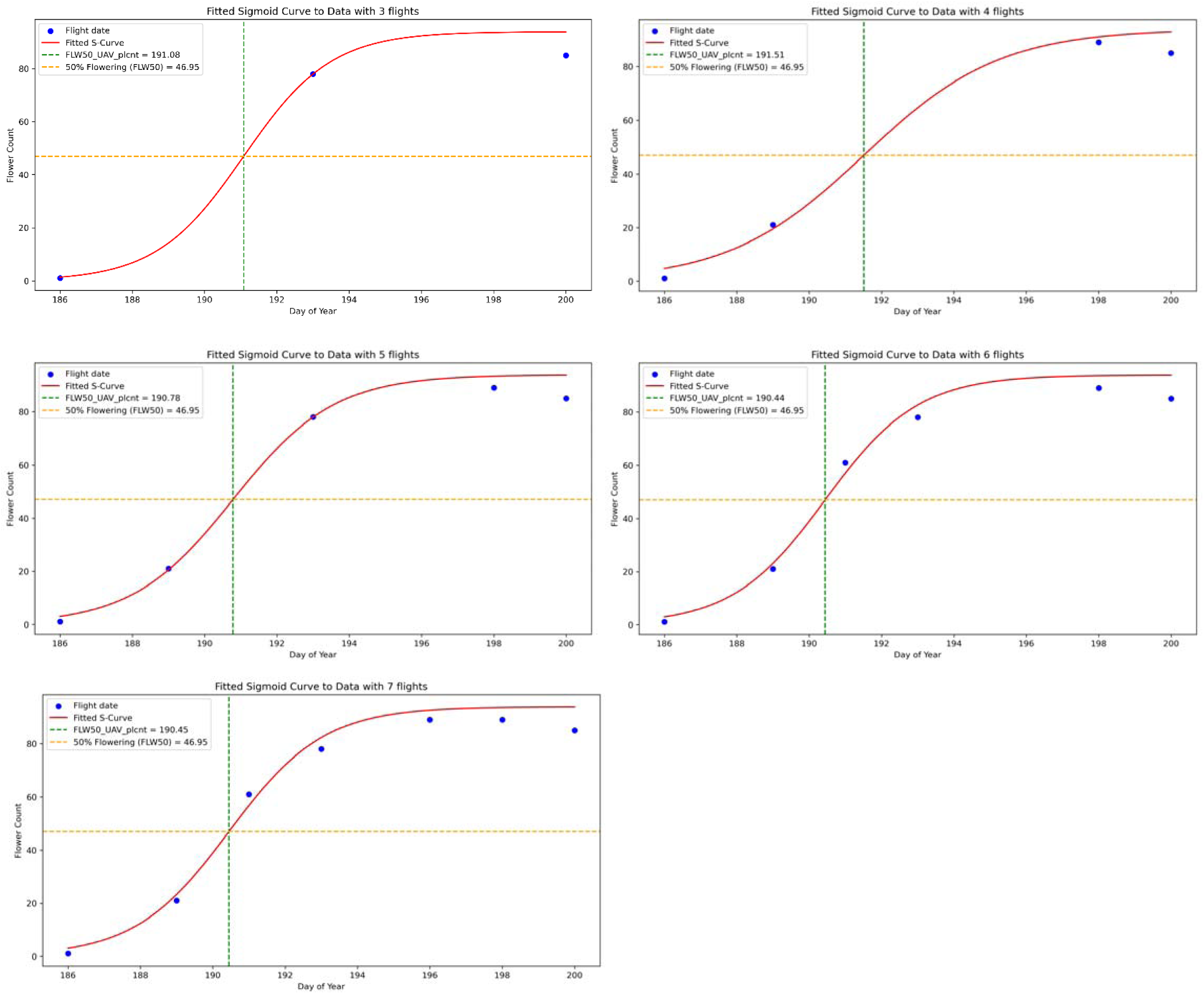
Fitted sigmoid curve detailing a visual representation of the data obtained using the various flight frequency configuration. A selected breeding plot was displayed to demonstrate the logistic growth trends. Blue dots represent the observed data points for flower counts obtained on specific days of the year during UAV flights. The red curve (Fitted S-curve) is the sigmoid curve fitted to the data points that represents the model’s estimation of the relationship between the day of the year and the flower count. The sigmoid shape indicates that the flower count increases with time, approaching an upper limit as time progresses. The Horizontal Dashed Orange Line represents the flower count that corresponds to the estimated 50% of the maximum flower count predicted by the model. Vertical Dashed Green Line indicates the model’s estimated midpoint, the point at which 50% of the maximum flower count is reached.

**Supplemental Figure 3:**
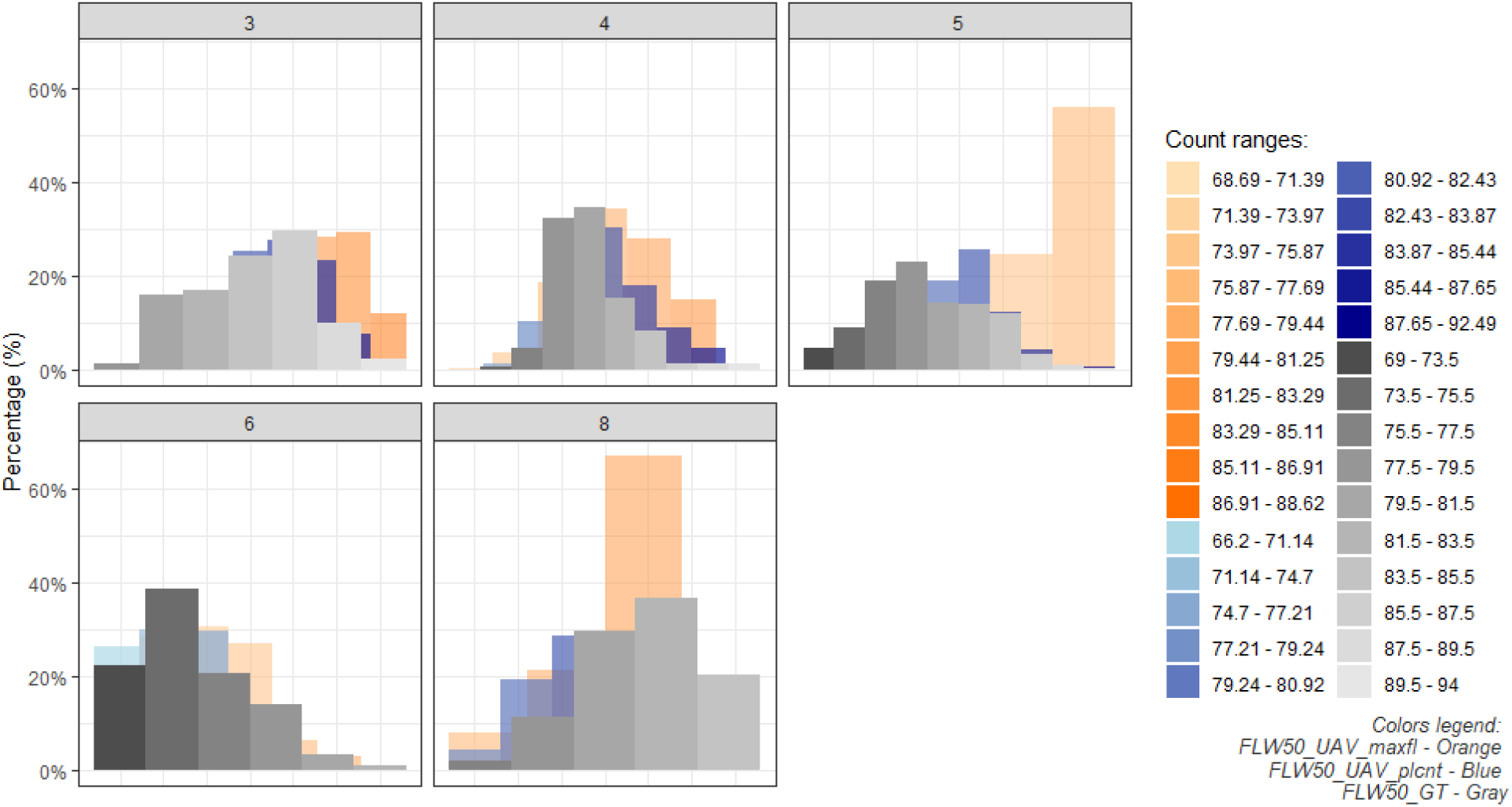
Frequency Distribution of 50% Flowering Dates Across Selected Trials. The histograms represent the percentage of observations falling within specific flowering date ranges for ground truth (FLW50_GT - Gray), UAV-derived plant stand counts (FLW50_UAV_plcnt - Blue), and UAV-derived head flower detections (FLW50_UAV_maxfl - Orange) for trials 3, 4, 5, 6, and 8. The color-coded count ranges key aids in distinguishing the distribution patterns of the flowering dates observed in each method.

**Supplemental Figure 4 a-b:**
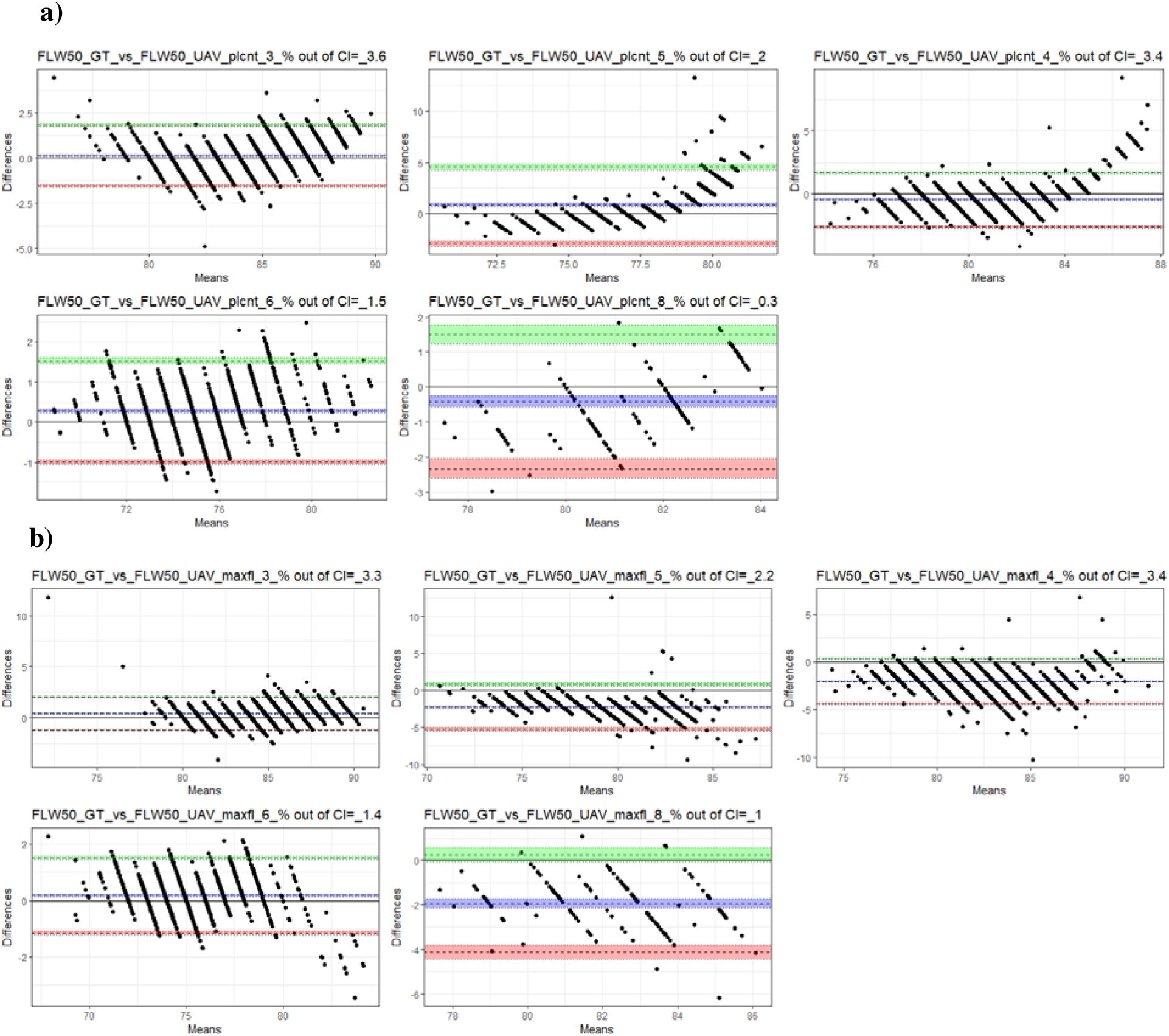
Bland-Altman Plots Assessing Agreement Between UAV-Derived Estimates and Ground Truth Flowering Dates Across Trials. The plots illustrate the mean differences and limits of agreement for each trial, calculated using ±1.96 for the 95% confidence intervals, highlighting the consistency of UAV-derived data (a - FLW50_UAV_plcnt, b - FLW50_UAV_maxfl) with ground truth (FLW50_GT). The trial identification and the percentage of observations outside the confidence interval (CI) are displayed at the top of each trial plot.

**Supplemental Figure 5:**
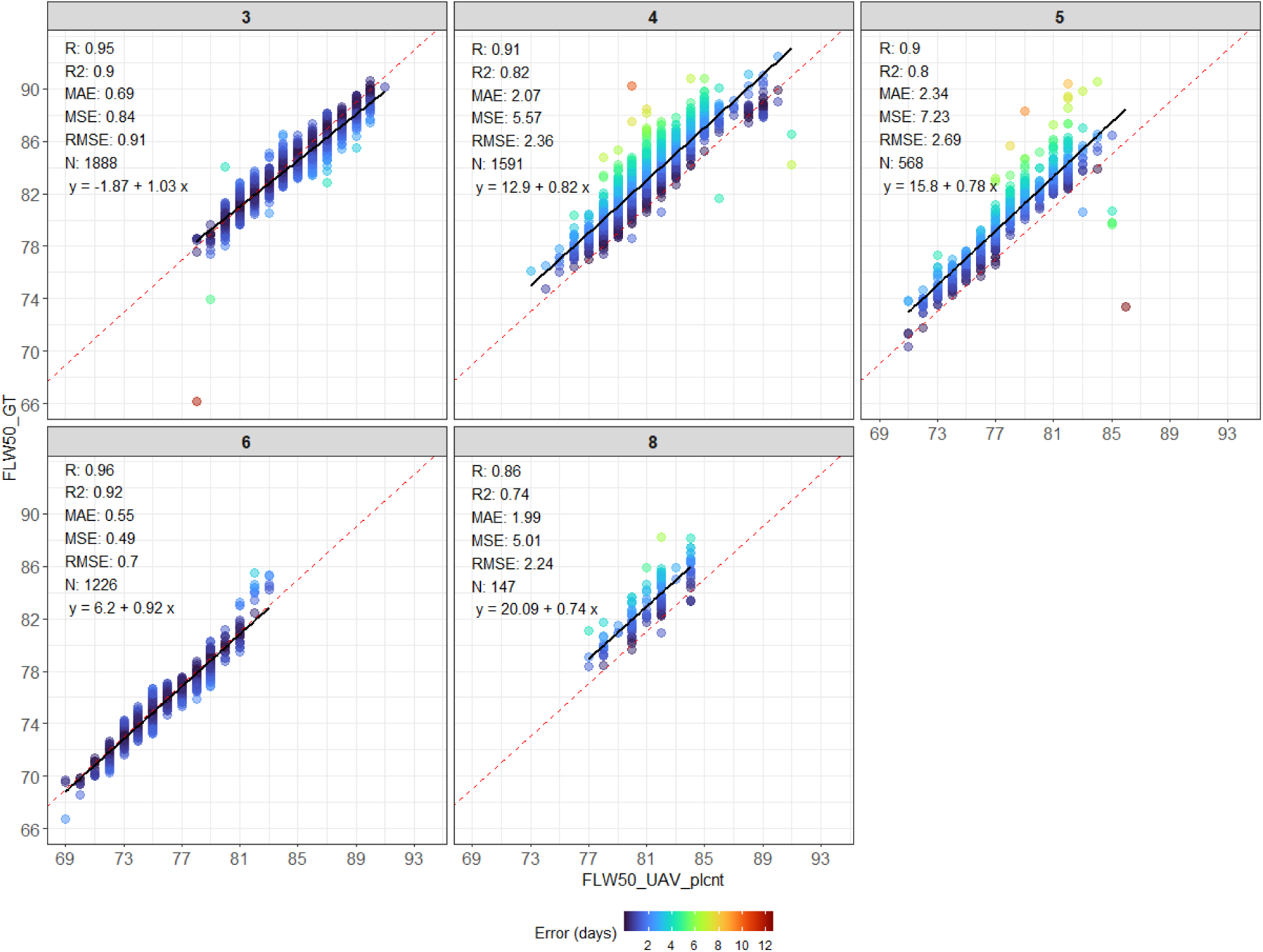
Correlation and error analysis of 50% flowering date estimations for sunflower trials for the validation data set. The scatter plots illustrate the relationship between the ground truth 50% flowering date (FLW50_GT) and UAV-derived estimates using plant stand counts (FLW50_UAV_plcnt) across 8 different trials. The color gradient indicates error magnitude in days.

**Supplemental Figure 6:**
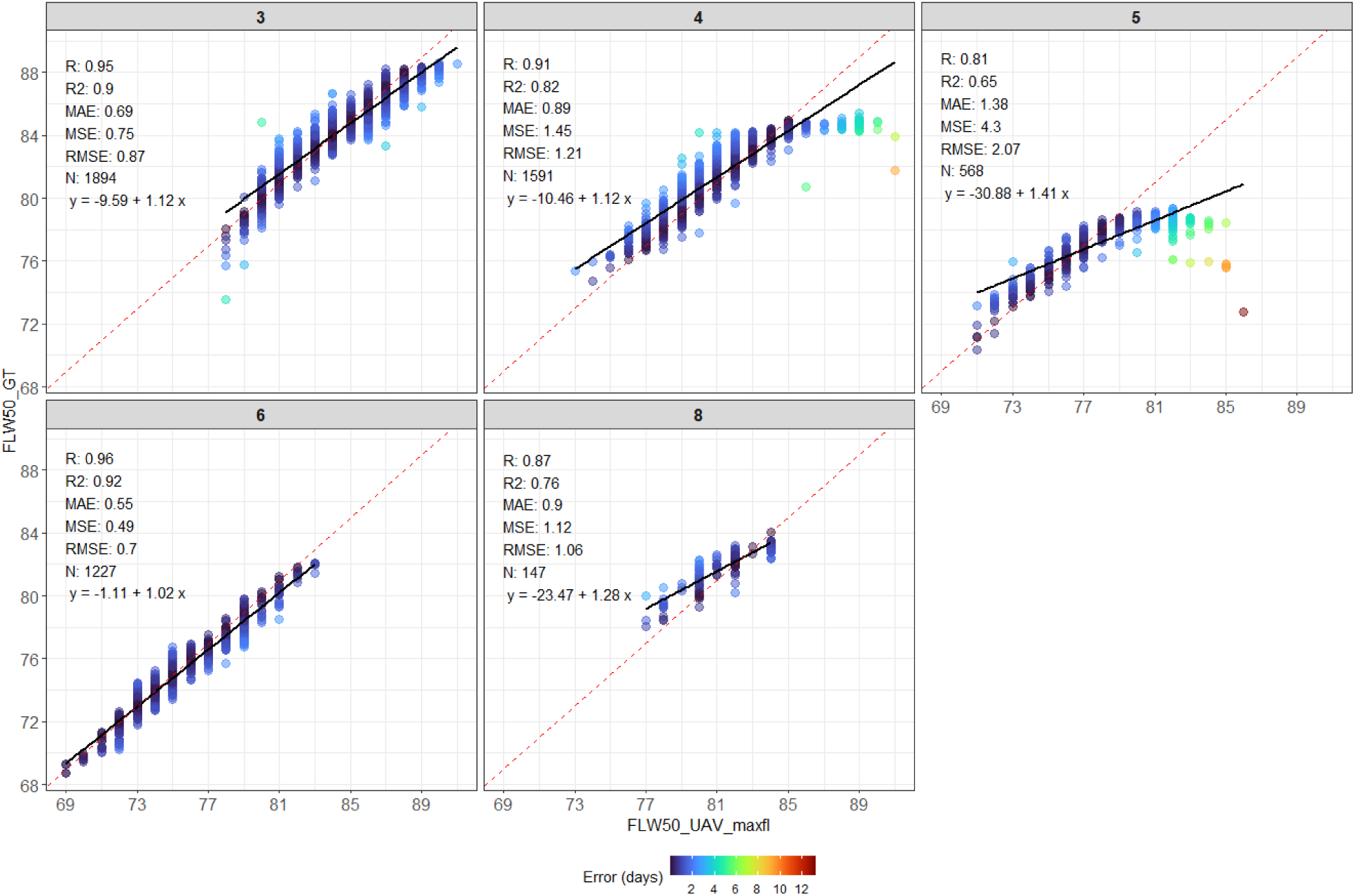
Correlation and error analysis of 50% flowering date estimations for sunflower trials for the validation data set. The scatter plots illustrate the relationship between the ground truth 50% flowering date (FLW50_GT) and UAV-derived estimates using maximum detected plants (FLW50_UAV_maxfl) across 8 different trials.

**Supplementary Figure 7:**
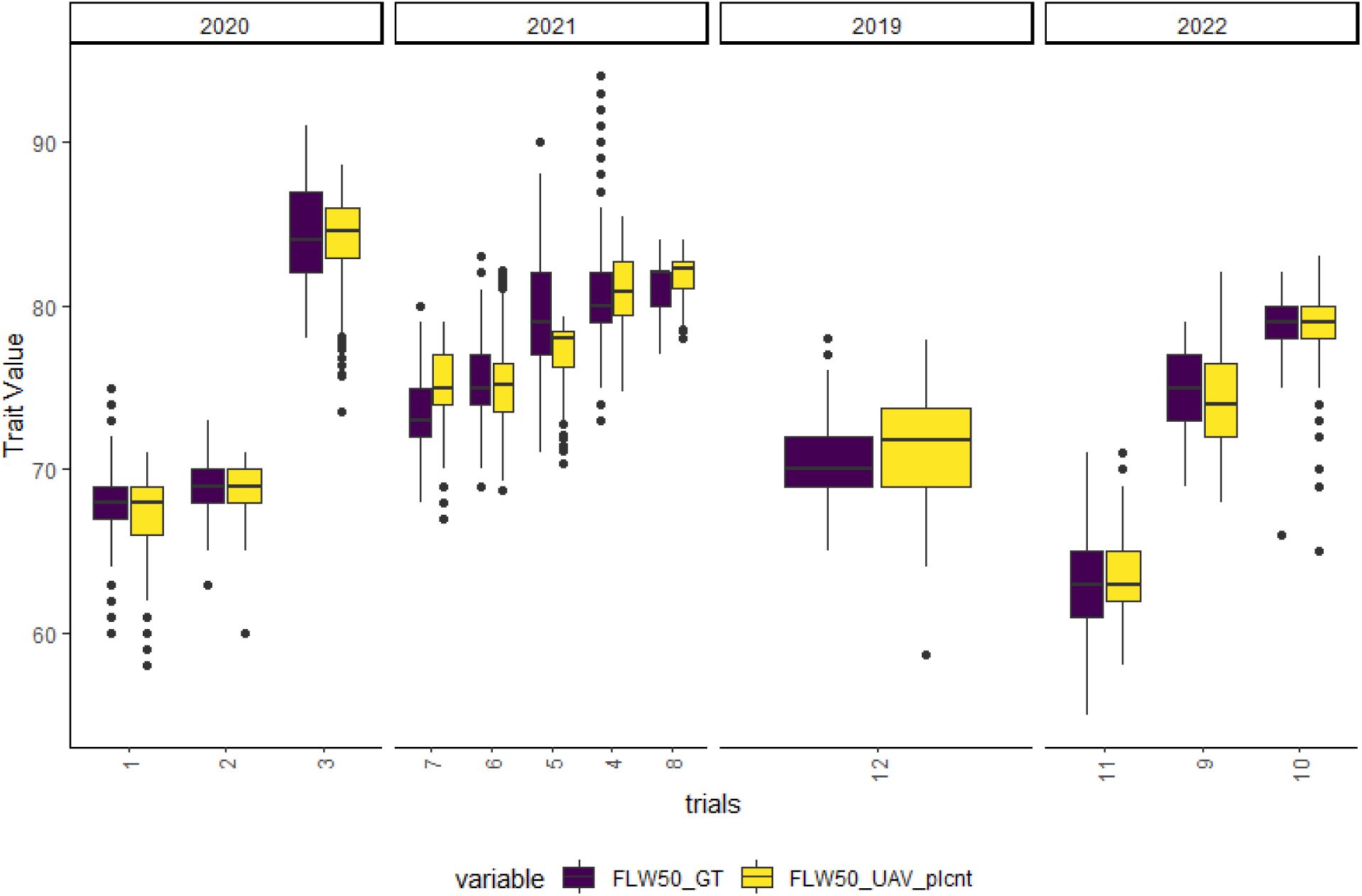
Comparative Raw Data Distribution of 50% Flowering Dates Across Trials. The boxplots depict both the GT dates (FLW50_GT) and UAV-derived plant stand count estimates (FLW50_UAV_plcnt), highlighting the data’s spread and central tendencies for each trial conducted in this study from 2019 to 2022.

**Supplemental Figure 8:**
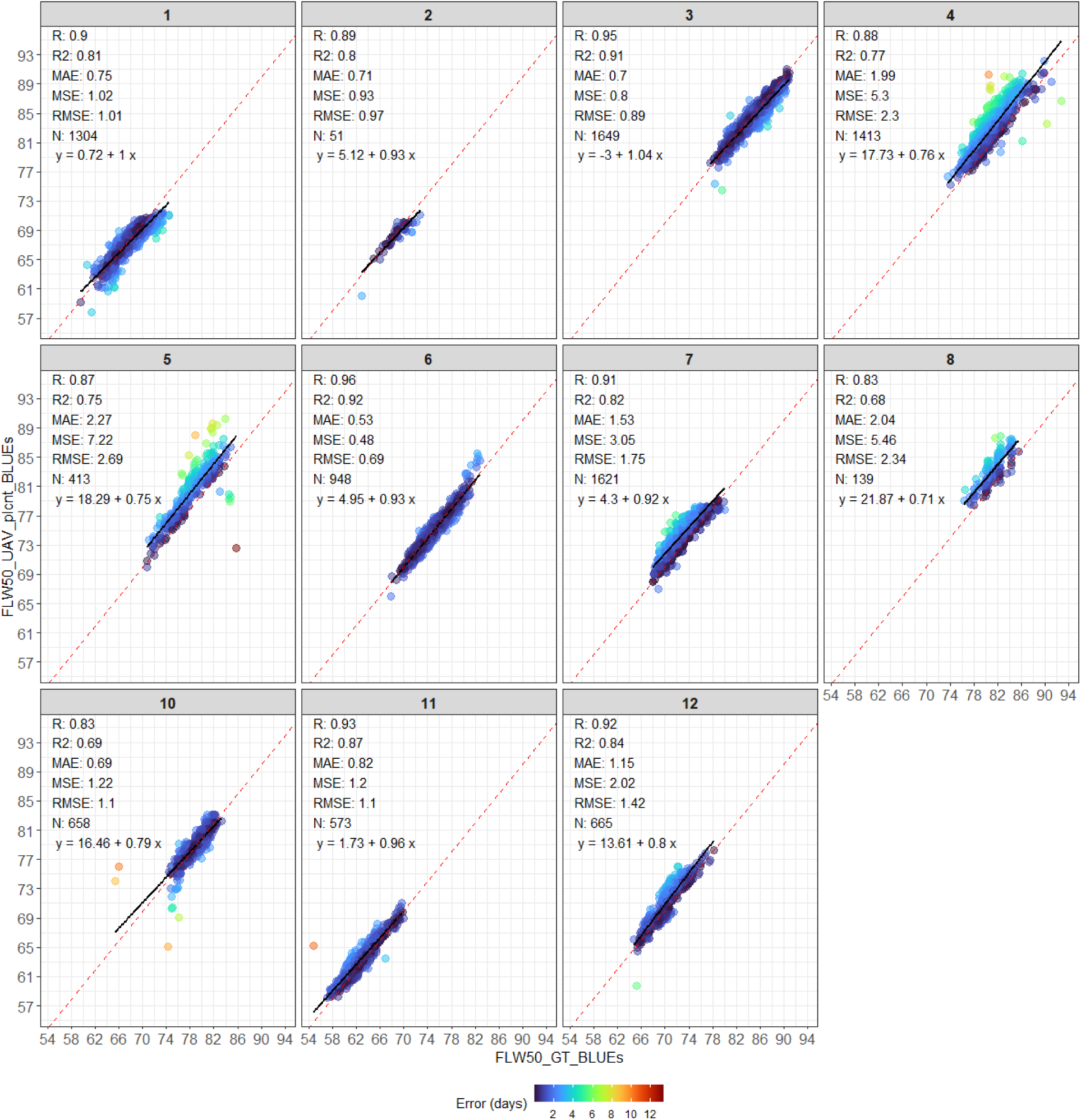
Correlation and error analysis of 50% flowering date estimations for sunflower trials using BLUEs estimation. The scatter plots illustrate the relationship between the ground truth 50% flowering date (FLW50_GT from BLUEs) and UAV-derived estimates using plant stand counts (FLW50_UAV_plcnt from BLUEs) across 11 different trials. The color gradient indicates error magnitude in days. Statistical metrics such as correlation coefficient (*R*), coefficient of determination (*R²*), mean absolute error (*MAE*), mean squared error (*MSE*), root mean squared error (*RMSE*), and linear regression equations are provided for each trial, demonstrating the accuracy of UAV estimations against field data.

**Supplemental Table 1:** Estimated variance components and genetic parameters for ground truth 50% flowering date (FLW50_GT) and UAV-derived estimates using plant stand counts (FLW50_UAV_plcnt) across 11 different trials from 2020 to 2022.

| FLW50_GT |  |  |  |  |  |  |
| --- | --- | --- | --- | --- | --- | --- |
| **Trials** | $\sigma_G^2$ | $\sigma_P^2$ | $\sigma_\varepsilon^2$ | $h^2$ | $h_g^2$ | $r_{gg}$ |
| **1** | 3.97<sup>**</sup> (81.90)<sup>§</sup> | 4.84 | 0.53 | 0.88 | 0.87 | 0.94 |
| **2** | 2.82<sup>**</sup> (81.93) | 3.44 | 0.56 | 0.83 | 0.86 | 0.94 |
| **3** | 3.79<sup>**</sup> (69.13) | 5.49 | 1.28 | 0.75 | 0.74 | 0.86 |
| **4** | 4.00<sup>**</sup> (63.57) | 6.3 | 1.79 | 0.69 | 0.68 | 0.83 |
| **5** | 12.96<sup>**</sup> (94.12) | 13.77 | 0.73 | 0.95 | 0.95 | 0.97 |
| **6** | 3.36<sup>**</sup> (66.25) | 5.07 | 1.18 | 0.74 | 0.73 | 0.87 |
| **7** | 3.78<sup>**</sup> (83.98) | 4.5 | 0.60 | 0.86 | 0.86 | 0.93 |
| **8** | 2.8<sup>*†</sup> (75.75) | 3.7 | 0.36 | 0.89 | 0.84 | 0.92 |
| **10** | 1.94<sup>**</sup> (71.78) | 2.7 | 0.76 | 0.72 | 0.72 | 0.87 |
| **11** | 5.31<sup>**</sup> (85.78) | 6.19 | 0.70 | 0.88 | 0.88 | 0.94 |
| **12** | 4.98<sup>**</sup> (91.27) | 5.46 | 0.39 | 0.93 | 0.92 | 0.96 |
| FLW50_UAV_plcnt |  |  |  |  |  |  |
| **1** | 3.07<sup>**</sup> (81.24) | 3.77 | 0.50 | 0.86 | 0.85 | 0.93 |
| **2** | 3.10<sup>**</sup> (93.22) | 3.33 | 0.21 | 0.94 | 0.95 | 0.97 |
| **3** | 3.09<sup>**</sup> (63.30) | 4.66 | 1.23 | 0.72 | 0.70 | 0.85 |
| **4** | 4.84<sup>**</sup> (75.53) | 6.67 | 1.14 | 0.81 | 0.79 | 0.99 |
| **5** | 10.78<sup>**</sup> (92.83) | 11.61 | 0.81 | 0.93 | 0.93 | 0.97 |
| **6** | 4.35<sup>**</sup> (78.49) | 5.54 | 0.77 | 0.85 | 0.84 | 0.92 |
| **7** | 3.77<sup>**</sup> (89.57) | 4.21 | 0.33 | 0.92 | 0.91 | 0.96 |
| **8** | 2.69<sup>**†</sup> (91.14) | 2.95 | 0.00 | 1.00 | 0.94 | 0.97 |
| **10** | 2.86<sup>**</sup> (86.17) | 3.31 | 0.44 | 0.87 | 0.86 | 0.94 |
| **11** | 5.33<sup>**</sup> (89.28) | 5.97 | 0.38 | 0.93 | 0.92 | 0.96 |
| **12** | 6.64<sup>**</sup> (88.33) | 7.52 | 0.68 | 0.91 | 0.91 | 0.95 |

$\sigma_G^2$ , genotypic variance; $\sigma_\varepsilon^2$ , residual variance; $\sigma_P^2$ , phenotypic variance; $h_g^2$ , Cullis broad-sense heritability; $h^2$ , broad-sense animal breeding heritability; $r_{gg}$ accuracy of genotype selection; $r_{ge}$ correlation between genotypic values across environments. \*\* Significant at $P < 0.001$ , and \* Significant at $P < 0.05$ ; <sup>†</sup> Variance components obtained using an alternative mixed model without spatial correction structure. <sup>§</sup> Parenthetical values indicate the percentage (%) of the observed phenotypic variance $\sigma_P^2$ .

